# Evaluation of nutrient stoichiometric relationships amongst ecosystem compartments of a subtropical treatment wetland. Do we have “Redfield Wetlands”?

**DOI:** 10.1101/220186

**Authors:** Paul Julian, Stefan Gerber, Rupesh K Bhomia, Jill King, Todd Z. Osborne, Alan L. Wright, Matthew Powers, Jacob Dombrowski

**Affiliations:** University of Florida, Soil and Water Sciences Department, Ft. Pierce, FL 34945; University of Florida, Soil and Water Sciences Department, Gainesville, FL, 32611; South Florida Water Management District, Water Quality Treatment Technologies, West Palm Beach, FL, 33406; University of Florida, Whitney Laboratory for Marine Bioscience, St Augustine, FL 32080

**Keywords:** decomposition, mineralization, Everglades, treatment wetlands

## Abstract

**Background:** Evaluation of carbon (C), nitrogen (N) and phosphorus (P) ratios in aquatic and terrestrial ecosystems can advance our understanding of biological processes, nutrient cycling and the fate of organic matter (OM) in these ecosystems. Eutrophication of aquatic ecosystems can change the accumulation and decomposition of OM which can alter biogeochemical cycling and alter the base of the aquatic food web. This study investigated nutrient stoichiometry within and among wetland ecosystem compartments (i.e. water column, flocculent, soil and above ground vegetation biomass) of two sub-tropical treatment wetlands with distinct vegetation communities. Two flow-ways (FWs) within the network of Everglades Stormwater Treatment Areas in south Florida (USA) were selected for this study. We evaluated nutrient stoichiometry of these to understand biogeochemical cycling and controls of nutrient removal in a treatment wetland within an ecological stoichiometry context.

**Results:** This study demonstrates that C, N, and P stoichiometry can be highly variable among ecosystem compartments and between FWs. Power law slopes of C, N and P within surface water floc, soil and vegetation were significantly different between and along FWs.

**Conclusions:** Assessment of wetland nutrient stoichiometry between and within ecosystem compartments suggests unconstrained stoichiometry related to P that conforms with the notion of P limitation in the ecosystem. Differences in N:P ratios between floc and soil suggest different pathways of organic nutrient accumulation and retention between FWs. Surface nutrient stoichiometry was highly variable and decoupled (or closed to decoupled, by our criteria), in particular with respect to P. We hypothesize that decoupling may be the imprint of variability in inflow nutrient stoichiometry. However, despite active biogeochemical cycles that could act to restore nutrient stoichiometry along the FW, there was little evidence that such balancing occurred, as the degree of stochiometric decoupling in the water column did change with distance downstream. This information is only the beginning of a larger journey to understand stoichiometric processes within wetland ecosystems and how it related to ecosystem function.

## Introduction

The study of nutrient stoichiometry, pioneered by Redfield (1934, 1958), laid the foundation of two important biogeochemical principles that later became basic tenets of ecological stoichiometry: (1) organisms have consistent carbon (C), nitrogen (N) and phosphorus (P) molar ratios and (2) the abundance of C, N and P in a system is regulated by interactions between organisms and their environment. These principles were supported by the similarity of measured N and P concentrations in marine plankton relative to the ratio of mineral forms of N (as nitrate [NO_3_]), P (as phosphate [PO_4_]) and non-calcite inorganic C in deep ocean water (Redfield 1934, 1958). The stoichiometric values of the Redfield ratio describe the average composition of marine organic matter (OM) and the requirements for remineralization of OM. Since its acceptance, the Redfield ratio has been debated and revisited frequently in light of new analytical methods, more data, and clarification of the frequent misrepresentations of the notable Redfield ratio (Lenton and Watson 2000; Geider and La Roche 2002). Despite this ongoing re-evaluation of the Redfield ratio most studies generally do not reject these conclusions but rather add subtlety and nuance (Sterner et al. 2008). Furthermore, the Redfield concept has been extended beyond marine ecosystems into freshwater and terrestrial ecosystems (i.e. lakes, streams, wetlands, forests, deserts, etc.) where C and nutrient concentrations are generally not part of a homogenous reservoir, are more variable between ecosystem compartments, residence times of nutrients in the system are shorter and biogeochemical dynamics differ significantly (Dodds et al. 2002; Dodds 2003; Cleveland and Liptzin 2007; Xu et al. 2013).

Prior studies have suggested that C:N:P ratios in soil are tightly constrained (i.e. abundance of nutrients is highly correlated) suggesting that for any given P concentration there is a comparable C or N concentration providing a Redfield-like stoichiometric ratio in both bulk soil and soil microbial biomass across forested, grassland and natural wetland ecosystems (Cleveland and Liptzin 2007; Xu et al. 2013). Redfield (1958) observed both the concentrations and ratio of elements in ocean water were constrained, indicating that, despite large variability in nutrient concentrations, the ratios remain unchanged in a specific ecosystem reservoir or compartment (i.e. water column, phytoplankton, sediment, etc.). Constrained stoichiometry suggests close interactions and feedbacks between organisms and their environment resulting in proportional scaling of nutrients (Fig 1). The relative ratios can change with abundance, suggesting a decoupling of nutrient cycles with increased nutrient availability. Here, we use coupling in a sense that there is an unbalanced stoichiometric budget expressed as disproportionate scaling between nutrients within a given ecosystem compartment, that may be allometric but predictable based on overall amount (high R^2^) or unpredictable (low R^2^). Shifts in nutrient concentrations within a given compartment can be driven by changes in nutrient loading, uptake and transport mechanisms or internal processes which in-turn can significantly alter stoichiometric composition of other ecosystem compartments resulting in an unbalanced stoichiometry cascade (Elser et al. 2009; Collins et al. 2017). Anthropogenically mediated nutrient loading to otherwise pristine aquatic ecosystems has potential to disrupt the ecological balance of nutrient supply and demand for both autotrophs and heterotrophs and can substantially affect nutrient (i.e. N and P) regeneration by disrupting productivity and nutrient remineralization. Long-term nutrient enrichment in aquatic ecosystems can affect overall nutrient abundance in all ecosystem compartments, affect rates of recycling between primary producers and OM decomposition, altering supply and demand for nutrients in different ecosystem compartments (plants, soil OM, microbial biomass) (Davis 1991; Reddy et al. 1999; Wright and Reddy 2001a). Therefore, excessive external inputs of nutrients to an ecosystem can potentially lead to a change of the stoichiometric balance of ecosystem compartments by preferential assimilation, changes in turnover and mineralization rates (Reddy and DeLaune 2008).

**Figure 1.**
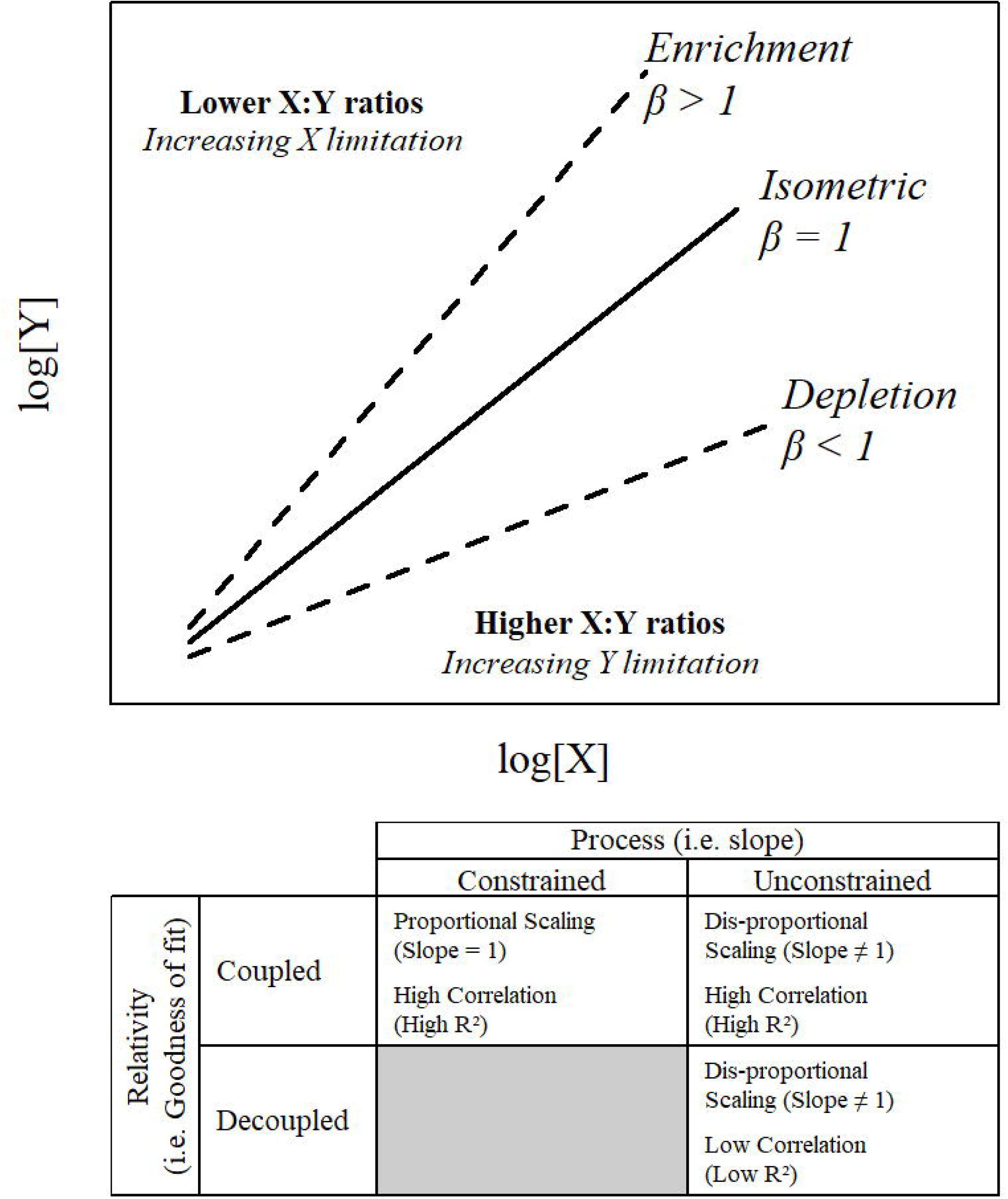
Top: Conceptual model for power law slope (β) interpretation relative to log transformed nutrient concentrations and relationship to stoichiometric ratios (i.e. X:Y). Bottom: Cross walk and function definitions of constrained/unconstrained and coupled/decoupled.

Treatment wetlands reflect an extreme end-member of such a disruption with long-term nutrient enrichment (Kadlec and Wallace 2009; Walker and Kadlec 2011). In an effort to restore the biological integrity of the Everglades ecosystem, the State of Florida and the US Federal government-initiated restoration and control efforts focusing on water quality improvement. One such effort is the construction and operation of constructed treatment wetlands to improve the quality of agricultural runoff water originating in the Everglades Agricultural Area (EAA) prior to entering the downstream Everglades ecosystem (Chen et al. 2015). These treatment wetlands, referred to as the Everglades stormwater treatment areas (STAs) were constructed with the primary objective of removing excess P from surface water prior to discharge to the Everglades Protection Area. The STAs are composed of several treatment cells or flow-ways (FWs) which use natural wetland communities to facilitate the removal of P from the water column by leveraging natural wetland processes including nutrient storage into vegetative tissues and burial within soils (Kadlec and Wallace 2009). The STAs are highly-managed treatment wetlands optimized to remove P by managing vegetation and regulating inflow and outflow volumes to optimize hydrologic residence times, hydraulic loading rates (HLR) and P loading rates (PLR) (Howard-Williams 1985; Kadlec and Wallace 2009).

In the Everglades STAs, water column P concentrations decline along the flow-way establishing a strong inflow-to-outflow nutrient gradient pointing to notable P sequestration (Juston and DeBusk 2011; Corstanje et al. 2016). This water column P gradient facilitated by long-term P loading has promoted the formation of a soil nutrient gradient spatially distributed from inflow-to-outflow (Zamorano et al. 2018). The strong P-gradient in surface water and soil compartments, suggest that other biologically relevant elements (C and N) may reflect this gradient to some degree (UF-WBL 2017). Gradients of C and N are apparent but vary in the degree of change along the FWs (UF-WBL 2017). Moreover, these treatment wetland ecosystems are typically heterogenous and exhibit significant differences in nutrient concentrations between ecosystems compartments (such as vegetation, floc and soils) as influenced by various wetland features and processes (Newman et al. 2004; Osborne et al. 2011b; Bhomia and Reddy 2018; Zamorano et al. 2018).

There is considerable variability in loading and storage depending on the location within a FW. Because other macro-elements (C and N) reflect to some degree changes in P may be possible to discern a Redfield-like ratio within STA FWs. What is not known is if this Redfield-like ratio is consistent among systems with different vegetation communities and along the FW with decreased nutrient concentration in the water column from inflow to outflow. Therefore, this study focuses on the evaluation of nutrient stoichiometric relationships within wetland ecosystem compartments along strong nutrient gradients and explore similarities or differences as a result of different vegetation. Our objective is to evaluate overall nutrient relationships (i.e. C x N, C x P and N x P) within surface water, soil flocculent material (floc), recently accreted soil (soil) and vegetation live aboveground biomass (AGB) between two FWs, one dominated by emergent aquatic vegetation (EAV) and the other by submerged aquatic vegetation (SAV). Using standard major axis (SMA) regression, it was tested as to whether nutrient concentrations scale allometric (independently) or isometric (dependently). The isometric model suggests a Redfield-like nutrient relationship that is strongly governed by a fixed biotic elemental ratio while allometric relationships imply shifts in nutrient ratios as concentrations (or amounts) of one nutrient changes. Differences in scaling relationships can indicate changes or differences in biogeochemical drivers and processes as suggested by Brown et al. (2002). We explored, how these relationships change along the FW of the treatment wetland (reflecting changes in nutrient supply) and between FWs (reflecting differences in vegetation). We first hypothesize that nutrient stoichiometric ratios in individual ecosystem compartments will differ between FWs with different vegetation types. Our second hypothesis is that the compartment’s stoichiometry do not follow the Redfield relationship (i.e. are *unconstrained*) but instead scale allometrically because of a large gradient in nutrient supply, and that the ratios change predictably and remain *coupled* (regression on SMA explains > 25 % of variability) within a FW because of the homogenous vegetation (i.e. the system’s macronutrients remain coupled). We further hypothesize that stoichiometric relationships will change along the flowway from inflow to outflow due to decreasing nutrient loading.

## Methods

### Study Area

A total of six STAs with an approximate area of 231 km^2^ are located south of Lake Okeechobee in the southern portion of the EAA (Fig 2). Prior land uses within the current STA boundaries include natural wetlands and agricultural land use dominated by sugarcane. The primary source of inflow water to the STAs is agricultural runoff originating from approximately 284 km^2^ of agricultural land use upstream. Everglades STA treatment cells are comprised of a mixture of EAV and SAV communities in several configurations including EAV and SAV treatment cells arranged in parallel or in series (Chen et al. 2015).

**Figure 2.**
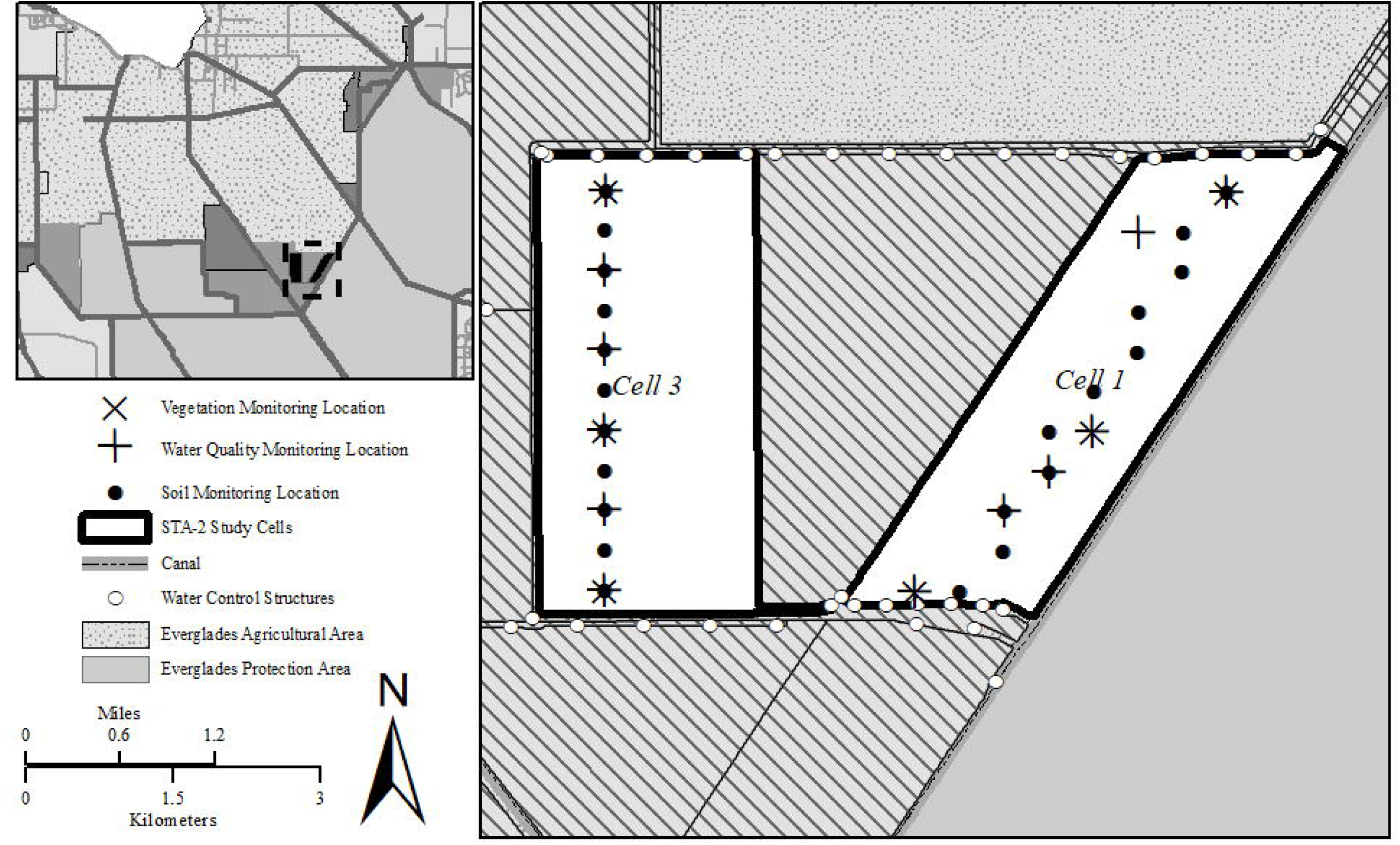
Surface water, soil and vegetation monitoring locations within Everglades Stormwater Treatment Area-2 Cells 1 (right) and 3 (left). Cell 1 is predominately emergent vegetation and Cell 3 is predominately submerged aquatic vegetation. Operationally these cells are identified as flow-way 1 and 3, respectively.

Stormwater Treatment Area-2 has been in operation since June 1999 with an effective treatment area of approximately 63 km^2^ divided into eight treatment cells. This study was conducted in two cells, FWs 1 and 3, respectively. The vegetation community of FW 1 is comprised predominately of EAV including *Typha domingensis* Pers. (cattail) and *Cladium jamaicense* Crantz (sawgrass) while FW 3 mainly consists of SAV including *Chara* spp. (muskgrass), *Potamogeton* spp. (pondweed) and *Najas guadalupensis* Spreng (southern naiad), periphyton communities typically in the lower two-thirds of the FW. Approximately a third of the FW is occupied by EAV species (Dombrowski et al. 2018). Furthermore, prior to STA-2 construction, FW 1 was a historic natural wetland while approximately two-thirds of FW 3 was previously farmed and is now managed as a SAV system (Juston and DeBusk 2006).

### Data Source

Data used in this study were collected by South Florida Water Management District and University of Florida and was a part of a larger project within the overall South Florida Water Management District’s (SFWMD) Restoration Strategies Science Plan to improve the understanding of mechanisms and factors that affect P treatment performance (SFWMD 2012). Data from one study of the Science Plan to evaluate P-sources, form, fluxes and transformation process in the STAs was used for this study and can be found in UF-WBL (2017). Water quality monitoring locations were established along two FWs within STA-2 along a transect running from inflow to outflow of the FW (Fig 2). Weekly surface water grab samples were collected at monitoring locations within FWs 1 and 3 to characterize changes in nutrient concentrations and availability during prescribed/semi-managed flow event. Flow events were planned as a short duration (fixed temporal window) during which hydraulic flows to the system were maintained within a pre-determined range, and extensive monitoring was undertaken to ascertain system’s response to the controlled flow regime. These prescribed flow events were scheduled and cycled through various flow/no-flow sequences for FWs 1 and 3.

Surface water grab samples were collected at a depth of 10-15 centimeters just under the water surface, approximately mid-water depth whenever feasible. Water column parameters such as total P (TP), total N (TN), and dissolved organic C (DOC) were analyzed for these samples. In the Everglades system, the organic C pool is predominately composed of the DOC fraction with low particulate OC concentrations (Julian et al. 2017), therefore the bias of not including particulates in the C analysis is small. Soil samples were collected along the flow transects twice during the dry and wet seasons between 2015 and 2016 using the push-core method consistent with prior wetland soil studies (Bruland et al. 2007; Osborne et al. 2011a; Newman et al. 2017). Samples were extruded from the soil core tube and partitioned into floc and soil. Floc was characterized as the suspended unconsolidated material on top of the consolidated soil. It was poured into a secondary sampling container, allowed to settle for 4 hours supernatant water was removed via aspiration and remaining floc material was collected and analyzed. The consolidated soil underneath the floc layer was segmented with the 0 - 5 cm interval retained for analyses. Floc and soil samples were analyzed for percent ash, TP, TN, and TC. Loss-on-ignition (LOI) was calculated using percent ash values subtracted by 100 percent. Live AGB were collected from dominant vegetation in FW 1 and FW 3 at the end of the 2015 (November 2015) and 2016 (September 2016) wet seasons. Additional details regarding floc and soil sampling is discussed by UF-WBL (2017). Vegetation sampling locations were located at inflow, mid and outflow regions of the FWs within close proximity to the surface water and soil monitoring locations (Fig 2). Vegetation samples were collected from four to eight randomly placed 0.25 m^2^ quadrats adjacent to the identified sampling locations. Dry homogenized vegetation samples were analyzed for TP, TN and TC content consistent with U.S. Environmental Protection Agency approved methods (Table S1). Surface water inflow volume and TP concentrations were retrieved from the South Florida Water Management District (SFWMD) online database (DBHYDRO; www.sfwmd.gov/dbhydro) for each FW between May 1^st^ 2014 and April 30^th^ 2018 to include periods prior to sampling for this study. For purposes of this data analysis and summary statistics, data reported as less than method detection limit (MDL; Table S1) were assigned a value of one-half the MDL, unless otherwise noted.

### Data Analysis

Hydraulic and P loading rates (HLR and PLR, respectively) were calculated based on methods by Kadlec and Wallace (2009). Weekly surface water grab TP samples were collected at inflow and outflow structures and used to estimate inflow and outflow P-load amounts. Phosphorus loading rates were estimated using the daily TP load divided by FW area. Hydraulic loading rates were estimated by dividing flow volume by FW area. Surface water nutrient concentrations were converted from mass of nutrient per volume concentration (i.e. mg L^−1^) to molar concentrations (i.e. mM). Soil and floc concentrations were converted from mass of nutrient per mass of soil (i.e. g kg^−1^) to per area (moles m^−2^) by multiplying the nutrient concentration (g kg^−1^) with bulk density (kg m^−3^) and depth (m), and dividing by the nutrient (i.e. C, N and P) atomic weight.:

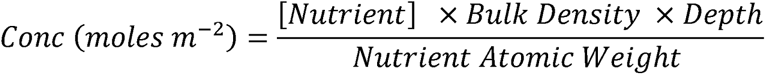

Nutrient concentrations in AGB was converted from mass per nutrient to mass of tissue to moles per area by multiplying nutrient concentration by biomass (g m^−2^) then dividing by the nutrient’s molecular weight. Expressing nutrient concentrations in moles per area units normalizes the concentration based upon bulk density for floc and soil and biomass in the case of AGB. In aquatic systems or in mineral soils Redfield ratios as well as evolution of isometric vs. allometric relationships have typically been carried out based on concentration (moles kg^−1^ or moles L^−1^). However, concentration-based analysis in the C-rich vegetation, floc and soil could be misleading, since any C increment would also increase the mass (the numerator for the concentration-based analysis; Fig S1). Hence, we chose to perform SMA analysis on a per area basis for these compartments.

Nutrient stoichiometric relationships within each ecosystem compartment (i.e. surface water, soil, floc and vegetation) were examined by evaluating power law slopes using standardized major axis (SMA) regression (’smatr’ package; Warton et al. 2006) consistent with Cleveland and Liptzin (2007). Unlike standard regression techniques which are used to predict one variable from another, SMA regression assesses the best fit line between two variables. Molar nutrient concentrations (water column) or amounts (mol m^−2^) were log-transformed and slope of the SMA regression was evaluated against the null hypothesis that the slope was not different from one (*β* ≠ 1). Power law (*y*= *kx ^β^*) and its linearized form (log(*y*) = *β* log(*x*) + log(*k*)) are used to evaluate the degree of proportional scaling between two variables. Scaling relationships and power-law distributions are key to understanding fundamental ecological relationships and processes in the natural system such as energy acquisition and transformation, biomass-growth relationships and evaluation of watershed chemostasis (Brown et al. 2002; Marquet et al. 2005; Wymore et al. 2017). In this analysis, we tested if the slope of the SMA regression results were statistically significantly different from one (i.e. ρ < 0.05) and interpreted as the variables are independent and do not proportionally scale (i.e. allometric growth) where one nutrient can either be enriched or depleted relative to the other (Fig 1). If the slope was not statistically different from one (i.e. ρ > 0.05), then the variables exhibited proportional changes (i.e. isometric growth; Fig 1) resulting in a constrained stoichiometry between nutrients. A slope not different from one would indicate that for any given concentration of nutrient X (i.e. C, N, or P) a proportional concentration of nutrient Y (i.e. P, C or N) existed. The degree of scaling (i.e. slope test) was combined with an evaluation of the regression coefficient of determination (R^2^) which indicated the degree of predictability (i.e. one nutrient can be used to predict the other). Low R^2^ values reflected high stoichiometric variability suggesting a decoupling of nutrients while high R^2^ values reflected low stoichiometric variability indicating a degree of coupling between nutrients. For our purposes, decoupled stoichiometric relationships were defined as a relationship with an R^2^ less than 0.25, R^2^ greater than 0.25 suggested some degree of coupling.

Standardized major axis SMA regression was applied to surface water, floc and soil nutrient concentrations or amounts separately between FWs to evaluate the overall (entire FW) stoichiometric relationship of C (DOC in surface water, TC in floc, soil and vegetation) to P, C to N and N to P using the ‘sma’ function in the smatr R-library (Warton et al. 2012). To compare nutrient stoichiometric relationships along each FW, monitoring locations were spatially aggregated to represent the inflow region (<0.3 fractional distance between inflow and outflow), mid region (0.3 – 0.6 fractional distance) and outflow region (>0.6 fractional distance) with FW region being evaluated using SMA regression. The resulting models from this location analysis were referred to as local models. Slope values from each FW (overall) and FW region (local) were compared to evaluate if each region shared a similar slope using a maximum likelihood comparison of slopes consistent with Warton and Weber (2002) using the ‘slope.com’ function in the smatr R-library. Additionally, slope values of overall stoichiometric comparisons were also evaluated between FWs. Ecosystem compartment nutrient concentrations and amounts as well as molar ratios were compared between FWs by Kruskal-Wallis rank sum test. To characterize the relationship between floc and soil along the two-flow path transects, regional categories outlined above (i.e. inflow, mid and outflow) were considered with soil and floc TN:TP being compared between FWs and distance downstream categories by Kruskal-Wallis rank sum test and a post-hoc Dunn’s test of multiple comparisons (‘dunn.test’ in the dun.test R-library) for each FW, separately. Floc and soil TN:TP were also compared by Spearman’s rank sum correlation by flow path separately. All statistical operations were performed with R© (Ver 3.1.2, R Foundation for Statistical Computing, Vienna Austria), unless otherwise stated all statistical operations were performed using the base R library. The critical level of significance was set at α = 0.05. Unless otherwise stated, mean values were reported with together with standard errors (i.e. mean ± standard error).

## Results

A total of six prescribed/managed flow events occurred between August 10^th^, 2015 and July 31^st^, 2017 with events ranging from 35 to 63 days in FWs 1 and 3 within STA-2, during which water column data were collected. During the flow events, daily HLR ranged between 0 (no inflow) and 31.5 cm d^−1^ with FW 3 receiving a relatively higher mean HLR of 3.4 ± 0.3 cm d^−1^ (Mean ± SE), compared to 2.2 ± 0.4 cm d^−1^ in FW 1 Observed daily PLR values ranged from 0 (no loading) to 90.9 mg m^−2^ d^−1^ with FW 1 receiving a higher relative load a mean PLR of 3.4 ± 0.7 mg m^−2^ d^−1^. Meanwhile, FW 3 experienced a mean PLR of 2.1 ± 0.2 mg m^−2^ d^−1^ (complete summary of flow event characteristics can be found in the Supplemental Material; Table S2 and Fig S2). The daily HLR and PLR observed during this study was consistent with historic operational loading rates experienced by these FWs (Chen et al. 2015). Furthermore, this synoptic comparison is also consistent with the recent period of record (last four water years) where HLR and PLR were generally greater in FW 3 than FW 1 (Fig S3). Mean HLR values observed during the study occurred at 56% and 71%, respectively for FW 1 and FW 3 along the HLR cumulative distribution function curve (CDF; Fig S3). Meanwhile, mean PLR values observed during the study occurred at 71% for both FW 1 and FW 3 along their respective PLR cumulative duration curve despite FW 1 having a higher maximum PLR and steeper CDF curve (Fig S3).

### Water Column C:N:P

Dissolved organic carbon, TN and TP concentrations were significantly different between FWs. Flow-way mean DOC concentrations (χ^2^ = 66.2; df=1;ρ<0.01; Table 1) and TN concentrations χ^2^ = 121.9; df=1;ρ<0.01; Table 1) were significantly greater for FW 3 than FW 1. Meanwhile, FW mean TP concentrations were significantly greater for FW 1 than FW 3 (χ^2^ = 15.5; df=1;ρ<0.01; Table 1). These differences are surprising given that each FW receives identical sources of water and presumably due to different loading regimes (HLR and PLR; Table S1) and contain different dominant vegetative communities resulting in differences in overall biogeochemical cycling. Across FWs, surface water DOC:TP values range from 216 to 14,613 (on a molar basis) with FW 3 having significantly greater values (χ^2^= 38.6, df=1,ρ<0.01; Table 1). Meanwhile, surface water DOC:TN values range from 9.3 to 24.0 with FW 1 having significantly greater mean DOC:TN values (χ^2^= 88.3, df=1,ρ<0.01; Table 1). Stoichiometric ratios of TN:TP ranged from 15.5 to 788.7 across the FWs with FW 3 having significantly greater TN:TP values than FW1 (χ^2^= 58.7, df=1,ρ<0.01; Table 1).

**Table 1.**
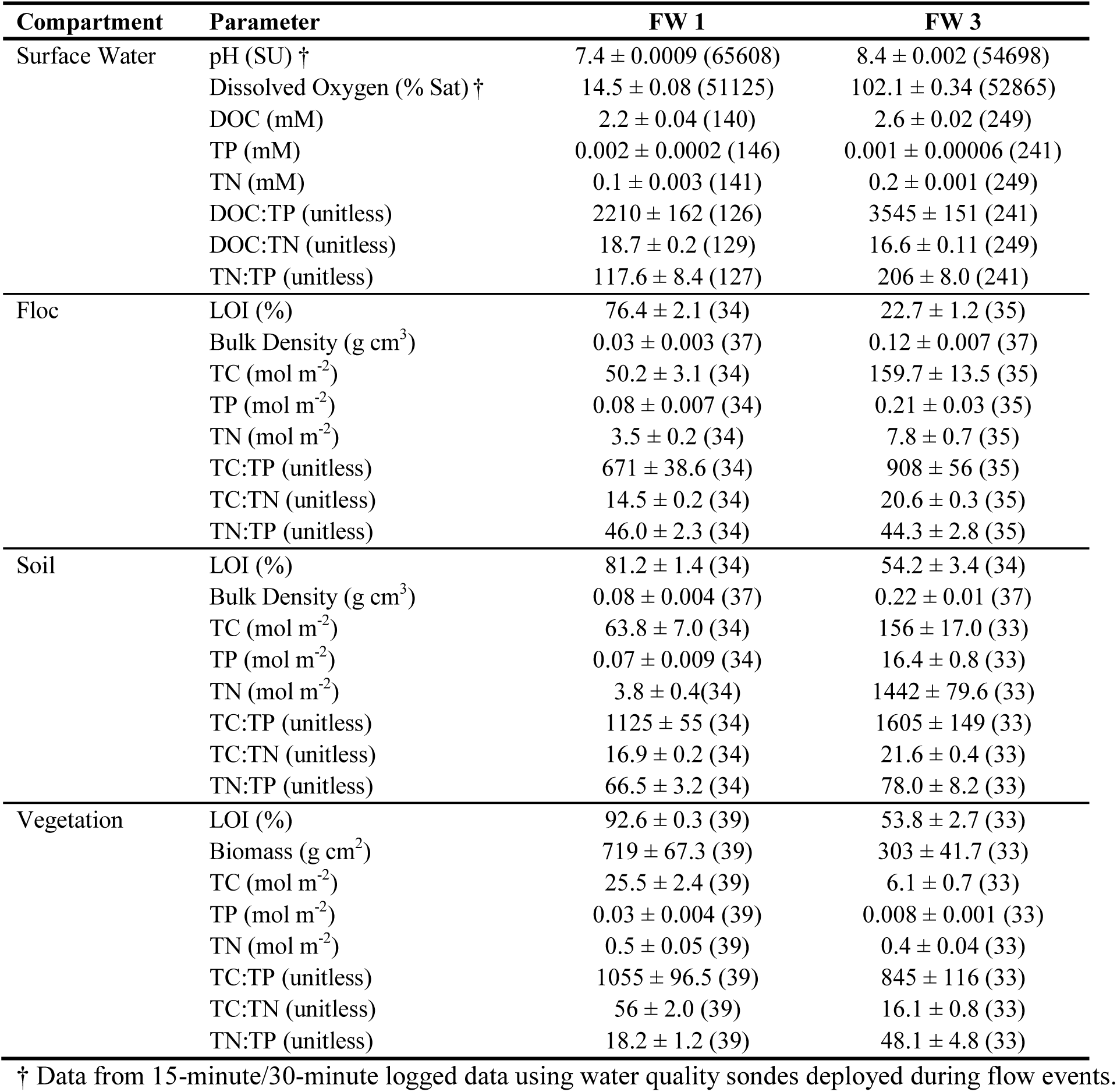
Summary statistics for parameters and matrices used in this study of samples collected along the Flow ways 1 and 3 flow-path transect within Stormwater Treatment Area-2. Summary statistics expressed as mean ± standard error (sample size). Matrices include surface water, soil flocculent material, recently accreted soil and living aboveground biomass of sampled vegetation. Stoichiometric ratios are expressed as molar ratios and are unitless. (DOC = Dissolved Organic Carbon; TP = Total Phosphorus; TN = Total Nitrogen; TC = Total Carbon; LOI = Loss-On-Ignition).

Overall FW surface water stoichiometric scaling relationships between DOC, TP and TN resulted in statistically significant relationships with slopes significantly different from one (Table 2) indicating that nutrient pools scaled independently (allometrically) in the surface water ecosystem compartment. Moreover, the R^2^ varies between models and FWs with the comparison of DOC to TP in FW 3 having a low R^2^ value (0.01) suggesting that the DOC to TP relationship was highly variable (use decoupled or unpredictable) for this flow-way. Overall FW model slopes were significantly different between FWs for comparisons of DOC to TP (Likelihood Ratio (LR) Statistics = 6.1, df=1, ρ<0.05), DOC to TN (LR Statistics = 5.0, df=1, ρ<0.05), and TN to TP (LR Statistics = 16.9, df=1, ρ<0.01).

**Table 2.** Surface water standardized major axis regression results for flow way 1 (FW 1) and FW 3 within Stormwater Treatment Area-2 for Inflow, Mid, Outflow and Entire FW (ALL) regions. Stoichiometric comparisons include Dissolved Organic Carbon to Total Phosphorus (DOC:TP), Dissolved Organic Carbon to Total Nitrogen (DOC:TN) and Total Nitrogen to Total Phosphorus (TN:TP).

| Compartment | Parameter | Region | FW 1 |  |  |  | FW 3 |  |  |  |
| --- | --- | --- | --- | --- | --- | --- | --- | --- | --- | --- |
|  |  |  | R<sup>2</sup> | Slope | F-value | p-value | R<sup>2</sup> | Slope | F-value | p-value |
| Water | DOC x TP | Inflow | 0.0001 | -0.24 | 155.8 | <0.01 | 0.0006 | -0.17 | 612.3 | <0.01 |
|  |  | Mid | 0.19 | 0.52 | 15.9 | <0.01 | 0.01 | -0.34 | 37.0 | <0.01 |
|  |  | Outflow | 0.22 | 0.31 | 150.2 | <0.01 | 0.02 | -0.26 | 437.3 | <0.01 |
|  |  | ALL | 0.23 | 0.23 | 705.6 | <0.01 | 0.01 | -0.18 | 1834.2 | <0.01 |
|  | DOC x TN | Inflow | 0.37 | 0.95 | 0.2 | 0.66 | 0.26 | 0.61 | 27.8 | <0.01 |
|  |  | Mid | 0.87 | 0.94 | 0.7 | 0.40 | 0.32 | 1.10 | 0.3 | 0.60 |
|  |  | Outflow | 0.91 | 0.81 | 28.5 | <0.01 | 0.57 | 1.17 | 8.6 | <0.01 |
|  |  | ALL | 0.78 | 0.77 | 38.7 | <0.01 | 0.34 | 0.90 | 4.3 | <0.01 |
|  | TN x TP | Inflow | 0.27 | 0.33 | 105.6 | <0.01 | 0.43 | 0.28 | 358.0 | <0.01 |
|  |  | Mid | 0.27 | 0.48 | 21.9 | <0.01 | 0.13 | 0.31 | 53.2 | <0.01 |
|  |  | Outflow | 0.21 | 0.35 | 107.4 | <0.01 | 0.08 | 0.22 | 672.8 | <0.01 |
|  |  | ALL | 0.41 | 0.28 | 569.7 | <0.01 | 0.28 | 0.20 | 2019.3 | <0.01 |

Much like the overall FW comparison of stoichiometric relationships, DOC by TP and TN by TP local (inflow, midflow and outflow) stoichiometric comparisons along the FWs resulted in models with slopes significantly different from one (Table 2) indicating that nutrient pools scaled independently (allometrically). With respect to DOC to TN, inflow and mid regions of FW 1 and the mid region of FW 3 resulted in models with slopes that were not significantly different from one demonstrating isometric scaling of nutrient pools (Table 2 and Fig 3). The remaining DOC by TN models had slopes significantly different from one and all models had relatively high R^2^ values along the inflow to outflow gradient, however it appears that R^2^ values were relatively lower at the inflow region of FW 1 and inflow and mid regions of FW 3 (Table 2). The R^2^ values of the TN by TP models remained relatively constant in FW 1 but declined along FW 3 indicating increased variability in the stoichiometric relationship of TN to TP in the SAV dominated FW. Stoichiometric relationships between DOC, TN and TP varied along both FWs (Fig 3). Slopes along FW 1 and FW 3 were significantly different for DOC to TP (LR Statistic=145.9, df=2, ρ<0.01 and LR Statistic=385.6, df=2, ρ<0.01, respectively). Slopes along FW 1 and FW 3 did not significantly differ for TN to TP (LR Statistic=3.3, df=2, ρ=0.19, LR Statistic =4.6, df=2, ρ=0.10, respectively) and DOC to TN slopes did not differ for DOC to TN (LR Statistic=3.8, df=2, ρ=0.15) along FW 1 indicating common slopes within each region of the FW. Meanwhile, slopes along FW 3 significantly differed for DOC to TN (LR Statistic=31.9, df=2, ρ<0.01).

**Figure 3.**
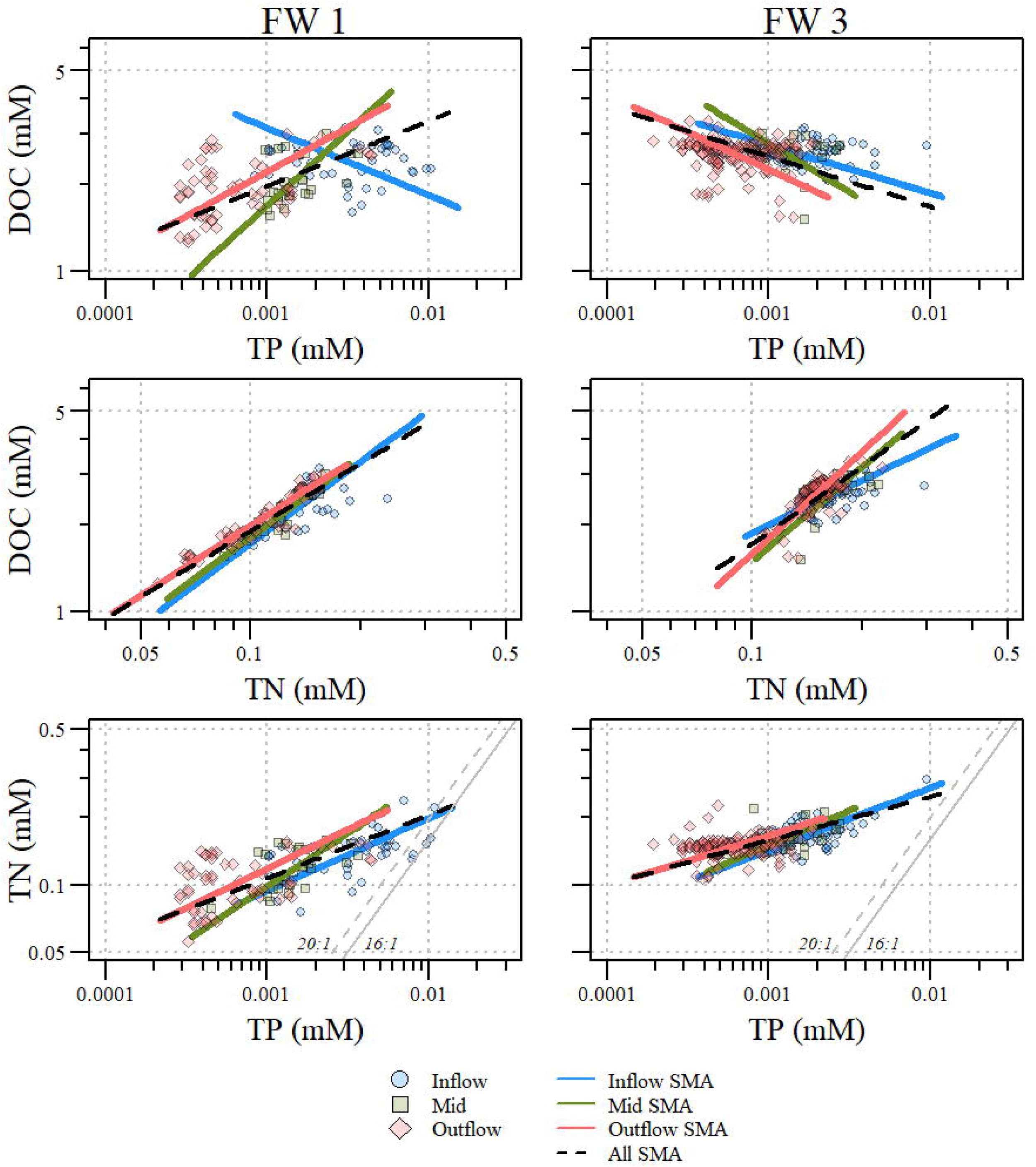
Stoichiometric relationships between dissolved organic carbon (DOC), total phosphorus (TP) and total nitrogen (TN) in the surface water ecosystem compartment for Stormwater Treatment Area 2, flow-ways (FWs) 1 and 3. Inflow, mid, outflow and overall standardized major axis (SMA) regressions are indicated by lines through the data. Values can be converted to mass per volume (i.e. milligram per liter) concentration by multiplying each value by its respective conversion factor (C = 12.01; N = 14.00; P = 30.97).

### Flocculent C:N:P

Floc in FW 3 had a greater mean bulk density and lower mean LOI than floc in FW 1, indicating that material in FW 1 is contains more organic material than FW 3 (Table 1). Much like the water column significant differences in floc nutrient content differed between FWs with FW 3 having significantly greater TP (χ^2^ = 18.9; df=1;ρ<0.01), TN (χ^2^ = 30.6; df=1;ρ<0.01) and TC (χ^2^= 44.5; df=1;ρ<0.01; Table 1) on an area basis (i.e. mol m^−2^). Across FWs, floc TC:TP values ranged from 411 to 1,369. Notable differences in stoichiometric ratios between FWs were apparent with C:P and C:N ratios being significantly different between FWs. Flow-way mean TC:TP and TC:TN values were significantly larger for FW 3 (χ^2^= 7.2, df=1,ρ<0.01, χ^2^= 46.7, df=1,ρ<0.01 respectively; Table 1). Meanwhile, floc TN:TP values did not significantly differ between FWs (χ^2^= 0.0006, df=1,ρ=0.94; Table 1). Comparisons of overall stoichiometric scaling between TC, TN and TP in the floc compartment resulted in slopes significantly different from one (allometric scaling) except for TC to TN comparisons in FW 3 with a reported slope of 0.99 indicating near isometry between TC to TN (Fig 4 and Table 3). The overall FW models for TC to TP and TN to TP, respectively, resulted in moderate R^2^ values for both FWs while TC to TN R^2^ values were much higher. Slopes of the overall FW models for TC to TP, TC to TN and TN to TP were not significantly different between FWs (LR Statistic = 0.08, df =1, ρ=0.77, LR Statistic = 1.35, df =1, ρ=0.25 and LR Statistic = 0.02, df =1, ρ=0.88, respectively).

**Figure 4.**
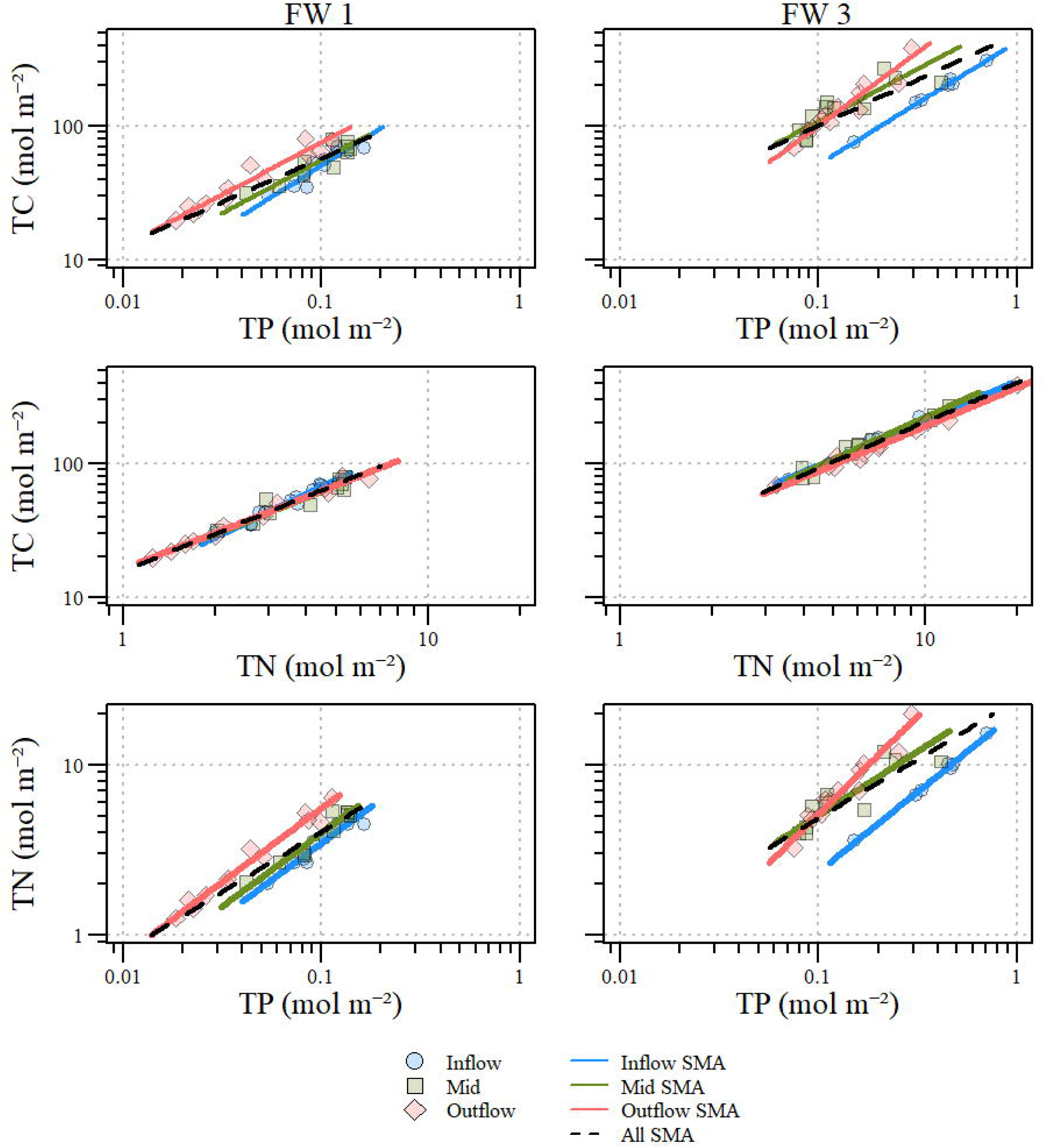
Stoichiometric relationships between total carbon (TC), total phosphorus (TP) and total nitrogen (TN) in the floc ecosystem compartment for Stormwater Treatment Area 2, flow-ways (FWs) 1 and 3. Inflow, mid, outflow and overall standardized major axis (SMA) regressions are indicated by lines through the data. Values can be converted to mass per volume (i.e. milligram per kilogram) concentration by multiplying each value by its respective conversion factor (C = 12.01; N = 14.00; P = 30.97).

**Table 3.**
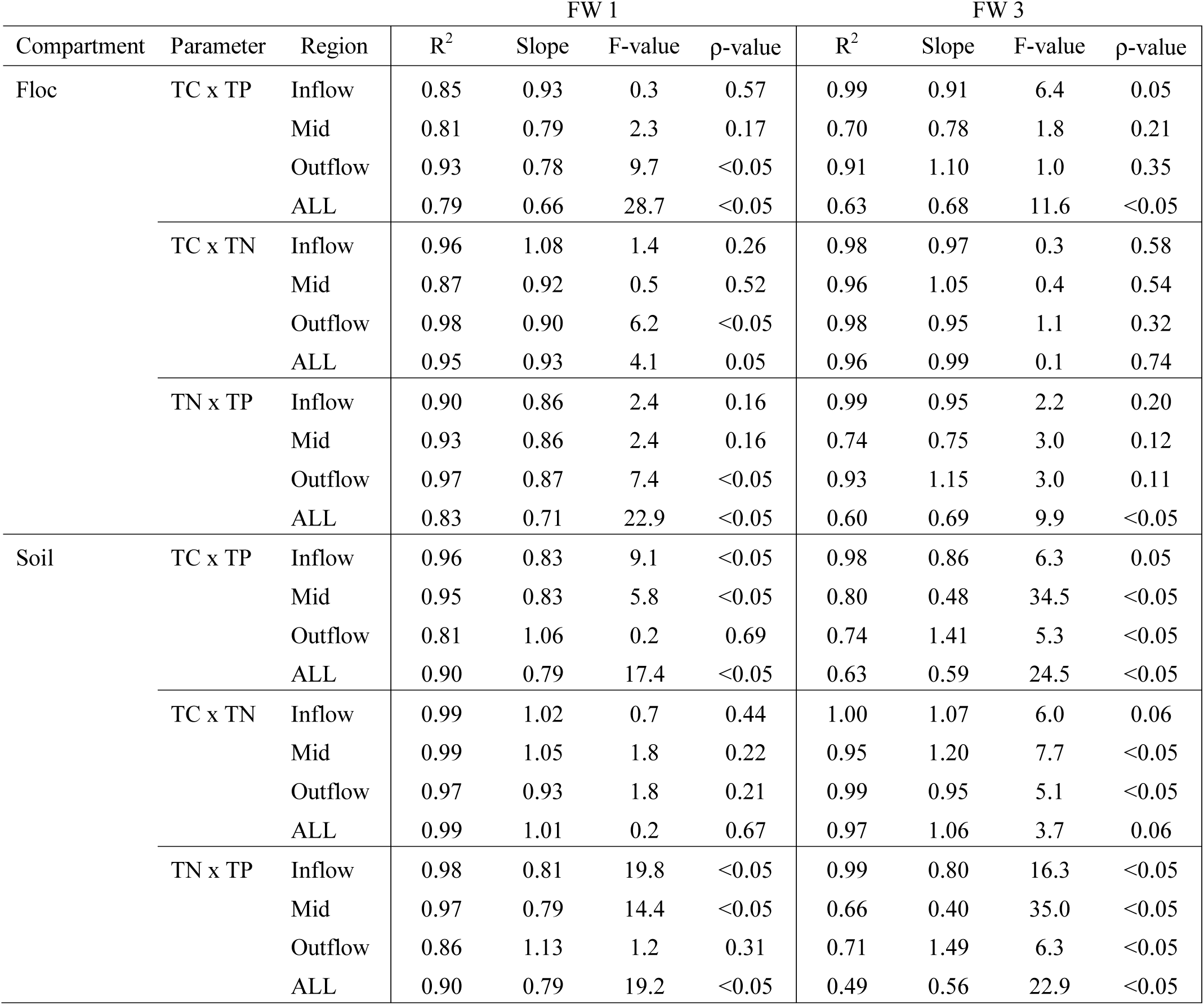
Flocculent and Soil standardized major axis regression results for flow way 1 (FW 1) and FW 3 within Stormwater Treatment Area-2 for Inflow, Mid, Outflow and Entire FW (ALL) regions. Stoichiometric comparisons include Total Carbon to Total Phosphorus (TC:TP), Total Carbon to Total Nitrogen (TC:TN) and Total Nitrogen to Total Phosphorus (TN:TP).

At a FW local scale (in-, mid, and outflow) in FW 1, TC to TP slopes at inflow and mid regions were not significantly different from one while outflow TC to TP was significantly different from one with a reported slope of 0.78 indicating potential depletion of C relative to P. Local TC to TP scaling relationships in FW 1 were highly variable with moderate R^2^ values at the inflow region and slightly higher R^2^ values at the outflow region indicating some stoichiometric variability despite slopes for inflow and mid regions not significant from one (Table 3). Total C to TP slope values in FW 3 along the FW (in-, mid and outflow) were not significantly different from one with some stoichiometric variability especially in the mid region (R^2^ = 0.70; Table 3). The TC to TN relationship for FW 1 outflow region was the only relationship with a slope significantly different from one with all other slopes being less than one indicating depletion of C relative to N (Table 3). Generally, local TC to TN R^2^ values were high except for mid regions of FW 1 with an R^2^ of 0.87 indicating some stoichiometric variability despite being an isometric relationship (Table 3). All local floc TN to TP slopes were not significantly different from one with FW 1 outflow region being the exception with a slope of 0.87 suggesting N depletion relative to P in this region of the FW (Table 3). All slopes along FW 3 with respect to TN to TP relationships were not significantly different from one (i.e. isometric scaling). A TN to TP slope of significantly different from one with a value of 0.32 for FW 3 mid region suggests possible depletion of N relative to P possibly marking a transition within the FW given the differences in TN to TP and TC to TP (Table 3). Stoichiometric relationships between floc TC, TN and TP varied along both FWs (Fig 4). Despite some variability in slope values along FW 1 and 3, slopes between regions were not significantly different for floc TC to TP (LR Statistic=1.41, df=2, ρ=0.49 and LR Statistic=3.49, df=2, ρ=0.17, respectively), TC to TN (LR Statistic=4.59, df=2, ρ=0.10 and LR Statistic=1.14, df=2, ρ=0.56, respectively) or TN to TP (LR Statistic=0.006, df=2, ρ=1.00 and LR Statistic=5.86, df=2, ρ=0.05, respectively).

### Soil C:N:P

Similar to the comparisons of floc nutrients, soil nutrient concentrations significantly differed between FWs with FW 1 having significantly greater TP (χ^2^ = 8.6; df=1;ρ<0.01), TN (χ^2^ = 17.1; df=1;ρ<0.01) and TC (χ^2^ = 26.3; df=1;ρ<0.01) concentrations than FW 3 (Table 1). Across FWs, soil TC:TP values ranged from 534 to 3551 and soil TC:TN ranged from 13.6 to 25.8 with significant differences for both ratios between FWs (χ^2^= 4.8, df=1,ρ<0.05 and χ^2^= 47.4, df=1,ρ<0.01, respectively). Both soil TC:TP and TC:TN values were higher for FW 3 than FW 1 (Table 1). Meanwhile, soil TN:TP values ranged from 22.3 to 179.4 with no significant difference between FWs (χ^2^= 0.6, df=1,ρ=0.45).

Comparisons of overall stoichiometric scaling relationships between TC, TN and TP in the soil ecosystem compartment resulted in slope values significantly different from one except for TC to TN comparisons indicating overall proportional (i.e. isometric) scaling of TC to TN in both FWs (Fig 5 and Table 3). Overall slope values for TC to TP and TN to TP comparisons in both FWs were less than one indicating depletion of C relative to P and depletion of N relative to P, respectively (Table 3). Coefficient of determinations (i.e. R^2^) for overall comparisons of TC, TN and TP varied between comparisons and FWs with TC to TN being very strongly coupled in both FWs while TC to TP and TN to TP are less strongly coupled in FW 3 as indicated by lower R^2^ values (Table 3). Overall FW model slopes were significantly different between FWs for comparisons of soil TC to TP (LR Statistics = 31.5, df=1, ρ<0.01) and TN to TP (LR Statistics = 31.6, df=1, ρ<0.01) while TC to TN were not significantly different between FWs (LR Statistics = 3.65 df=1, ρ=0.16).

**Figure 5.**
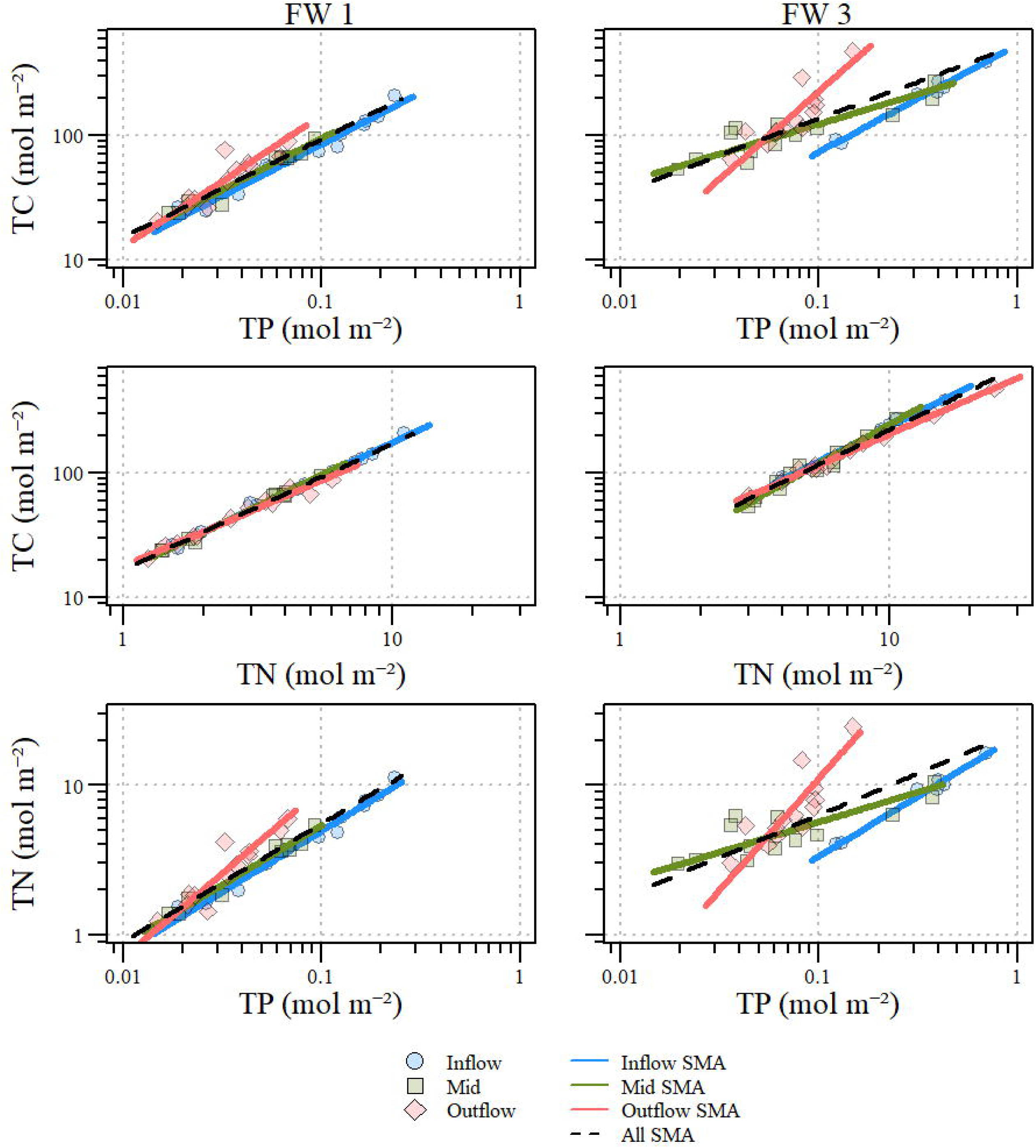
Stoichiometric relationships between total carbon (TC), total phosphorus (TP) and total nitrogen (TN) in the soil ecosystem compartment for Stormwater Treatment Area 2, flow-ways (FWs) 1 and 3. Inflow, mid, outflow and overall standardized major axis (SMA) regressions indicated by lines through the data. Values can be converted to mass per volume (i.e. milligram per kilogram) concentration by multiplying each value by its respective conversion factor (C = 12.01; N = 14.00; P = 30.97).

Local comparisons of soil TC, TN and TP resulted in a variety of scaling relationships both within and across FWs. Interestingly, within FW 1 TC to TP slopes were significantly different than one for inflow and mid regions with slope values less than one while outflow was not significantly different than one. This relationship was reverse in FW 3 where the inflow region was borderline significant (ρ-value=0.05) while mid and outflow slope values were significantly different than one moving from depletion to enrichment of C relative P (Table 3). Total C to TN relationships isometrically (i.e. proportionally) scaled for each region of FW 1, and the inflow region of FW 3 as indicated by slope values not significantly different from one (Table 3). However, TC to TN relationships for FW 3 mid and outflow relationships were significantly different from one with a slope value greater than one at the mid region suggesting enrichment of C relative to N and a slope value less than one at the outflow pointing to depletion of C relative to N (Table 3, Fig 5 and 9). Most of the soil TN to TP relationships were allometric (i.e. independently scaled) as indicated by slope values significantly different from one except for the outflow region of FW 1. For FW 1 and FW 3, inflow and mid regions had slopes less than one indicating the depletion of N relative to P. Meanwhile in the outflow region of FW 3, the TN to TP slope value greater than one indicated N enrichment relative to P (Table 3, Fig 5 and 9). All allometric TN to TP relationships had highly variable R^2^ values with the lowest values observed at the mid and outflow regions of FW 3, and none of the R^2^ values passed the decoupling threshold (i.e. 0.25). However, the relatively low R^2^ for FW 3 mid region suggested some degree of decoupling between TN and TP (Table 3). Scaling of variables along FW 1 (as indicated by slope values) were not significantly different with respect to TC to TP (LR Statistic = 2.4, df=2, ρ=0.30) and TC to TN (LR Statistic = 3.0, df=2, ρ=0.22) but were significantly different with respect to TN to TP (LR Statistic = 7.9, df=2, ρ<0.05). Alternatively, scaling of variables along FW 3 were significantly different for the comparison of TC to TP (LR Statistic = 18.5, df=2, ρ<0.01) and TC to TN (LR Statistic = 13.3, df=2, ρ<0.01), but not significantly different for scaling of TN to TP (LR Statistic = 3.6, df=2, ρ=0.16).

Further comparisons were made for floc and soil molar ratios as they are part of a decomposition continuum. Indeed, floc and soil N:P molar ratios were significantly correlated for sites within FW 3 (r_s_=0.80, _ρ_<0.01). In FW 1, it appears that soil TN:TP ratio has a maxima of 100 (with one exception), while there is a maxima in the floc compartment in FW 3, but not in the soil (Fig 6). Both floc (χ^2^ =12.7, df=2, _ρ_<0.01) and soil (χ^2^ =9.9, df=2, _ρ_<0.01) N:P molar ratios were significantly different along FW 3 with Floc N:P at the mid and outflow regions being statistically different from the inflow region (Z =-2.2, ρ<0.05 and Z=-3.6, ρ<0.01, respectively) while outflow and mid regions were similar (Z =-1.4, ρ=0.08). Similarly soil N:P values in FW 3 at the mid and outflow regions were statistically different from the inflow region (Z =-7.8, ρ<0.05 and Z=-3.1, ρ<0.01, respectively) while outflow and mid regions were similar (Z =-1.6, ρ=0.06). FW 1 soil and floc N:P values were positively correlated (r_s_= 0.64, ρ<0.01) and soil and floc N:P values were significantly different between FW regions (χ^2^ =12.0, df=2, _ρ_<0.01 and χ^2^=16.1, df=2, _ρ_<0.01, respectively). Floc N:P values in FW 1 were not significantly different between inflow and mid regions (Z=-1.6, ρ=0.06) but inflow (Z=-4.0, ρ<0.01) and mid (Z=-2.1, ρ<0.05) regions were significantly different from the outflow. However, soil N:P values were not significantly different between inflow and mid (Z=-1.6, ρ=0.054) or mid and outflow (Z=-1.6, ρ=0.054) but inflow was significantly different from outflow (Z=-3.5, ρ<0.01).

**Figure 6.**
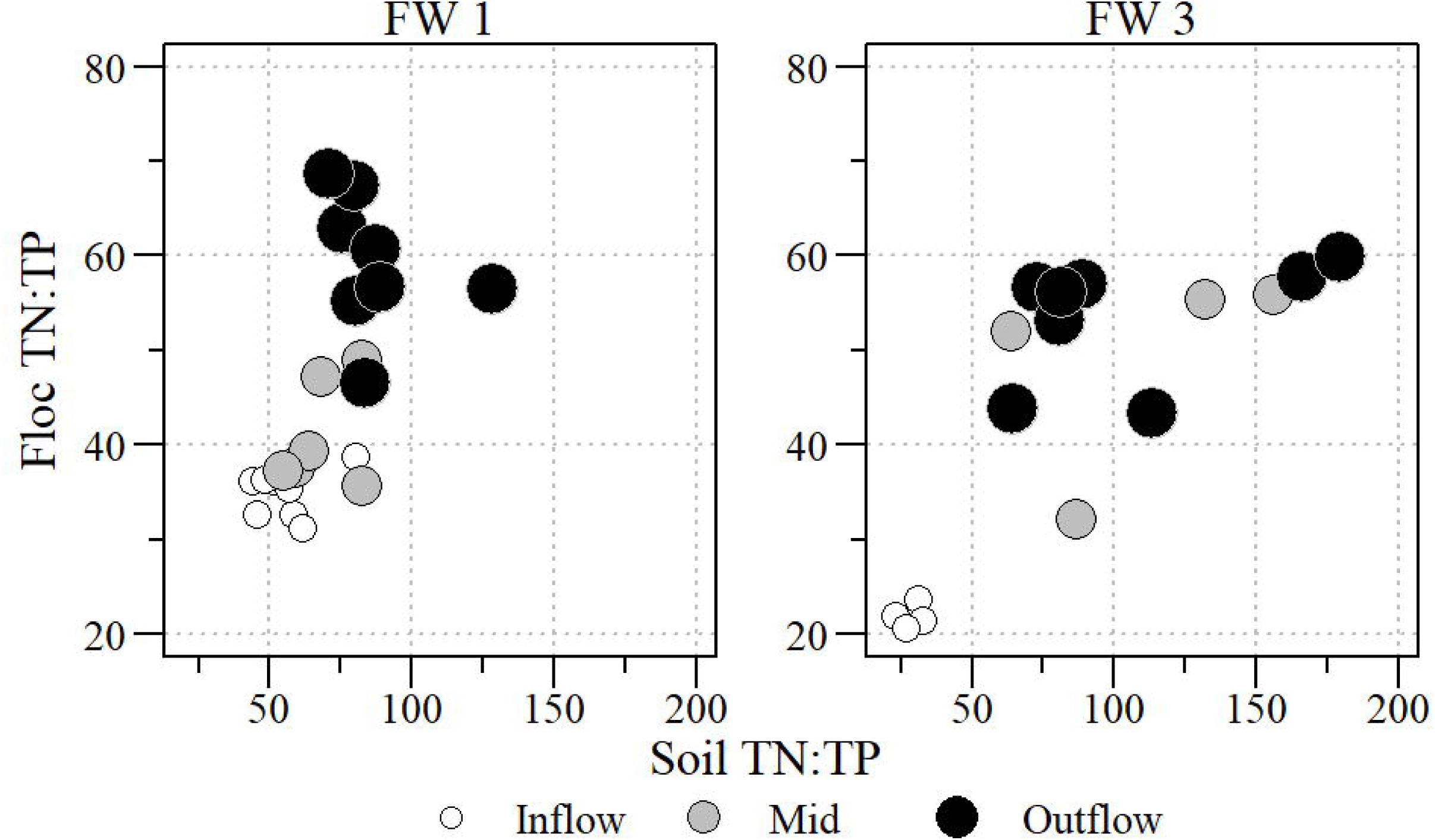
Comparison of floc and soil TN:TP molar ratio with location along the flow-way identified by size of point (i.e. larger point further down flow path) and distance categories.

### Vegetation C:N:P

Total P and TC concentrations on a per area basis (i.e. mol m^−2^) in live AGB were significantly different between FWs (χ^2^ = 30.0, df = 1, ρ<0.01 and χ^2^ = 40.1, df = 1, ρ<0.01, respectively) with FW 1 having higher TP and TC mass per area (Table 1). However, TN did not significantly differ between FWs (χ^2^ = 1.5, df = 1, ρ=0.22). Across FWs, live AGB TC:TP values ranged from 237.4 to 3,110 with FW 1 being significantly greater than FW 3 (χ^2^= 4.6, df=1,ρ<0.05; Table 1) due to the C rich EAV tissue. Live AGB TC:TN values ranged from 7.5 to 82.8 with FW 1 being significantly greater than FW 3 (χ^2^= 51.1, df=1,ρ<0.01; Table 1). Meanwhile, live AGB TN:TP value ranged from 7.4 to 120.4 with FW 3 being significantly greater than FW 1 (χ^2^= 42.2, df=1,ρ<0.01) (Table 1).

Stoichiometric comparison of live AGB nutrient concentrations varied across FWs. In FW 1, TC to TP and TC to TN comparisons resulted in slope values not significantly different from one indicating proportional (isometric) scaling of C to N and P in AGB tissue (Table 4). In FW 1, the TN to TP slope was significantly different from one with a slope less than one suggesting N depletion relative to P (Table 4). In FW 3, TC to TP and TC to TN comparisons resulted in slopes significantly different from one with slope values greater than one indicating enrichment of C relative to P and N in AGB tissue (Table 4). Unlike FW 1, the comparison of TN to TP in FW 3 resulted in slope not significantly different from one indicating isometric scaling of N and P in FW 3 AGB tissue (Table 4). Between all comparisons with slope values significantly different from one, R^2^ values were generally high ranging between 0.66 (FW 3; TC to TP) to 0.94 (FW 3; TC to TN Table 4) suggesting a high degree of coupling as expected. Between FWs, TC to TN slopes did not significantly differ (LR Statistic= 1.5, df =1, ρ=0.22) however it appears that intercept values significantly differ between SMA models driven by FW 1 having a greater C content (Fig 7). Slope values for TC to TP and TN to TP were significantly different between FWs (LR Statistic= 5.8, df =1, ρ<0.05 and LR Statistic= 5.4, df =1, ρ<0.05, respectively) where FW 3 had greater slope values for both comparisons (Table 4 and Fig 7).

**Figure 7.**
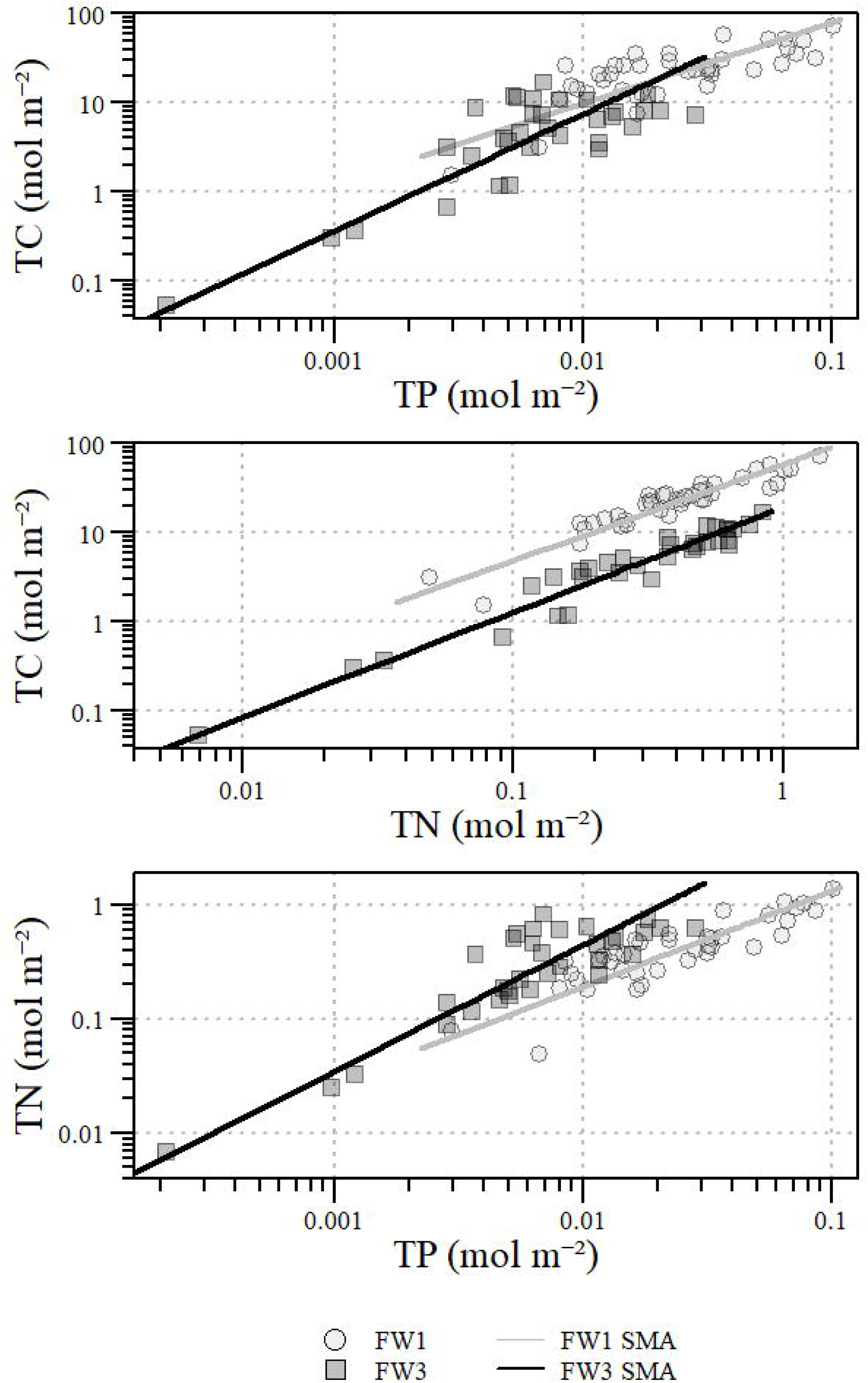
Stoichiometric relationships between total carbon (TC), total phosphorus (TP) and total nitrogen (TN) in vegetation ecosystem compartment for Stormwater Treatment Area 2, flow-ways (FWs) 1 and 3. Inflow, mid, outflow and overall standardized major axis (SMA) regressions indicated by lines through the data. Values can be converted to mass of nutrient per mass of soil (i.e. milligram per kilogram) concentration by multiplying each value by its respective conversion factor (C = 12.01; N = 14.00; P = 30.97).

**Table 4.**
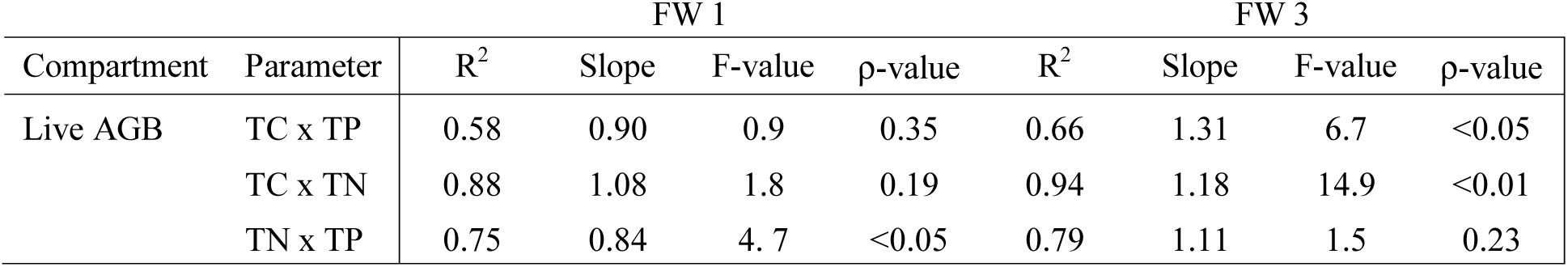
Live above ground biomass (AGB) nutrient concentration standardized major axis regression results for flow way 1 (FW 1) and FW 3 within Stormwater Treatment Area-2. Stoichiometric comparisons include Total Carbon to Total Phosphorus (TC:TP), Total Carbon to Total Nitrogen (TC:TN) and Total Nitrogen to Total Phosphorus (TN:TP).

## Discussion

The foundation of ecological stoichiometric theory is built on the laws of conservation of mass and constant proportions, which result in a long term balance of energy and elements in the context of interactions between ecosystem compartments (Helton et al. 2015; Van de Waal et al. 2018). This framework suggests a tight linkage between demand, use and recycling of nutrients which is ultimately the undercurrent of the Redfield ratio where nutrients are tightly constrained and driven by the interaction between the ambient environment and biota (Redfield 1958; Sterner and Elser 2002). Redfield (1958) observed that the abundance and ratio of elements in oceanic systems are constrained leading to the conclusion that a close interaction between organisms and internal biogeochemical processes regulate the similarities between the environment and the organisms manifesting in the characteristic “Redfield ratio”.

The predictive power of the Redfield ratio has prompted ecologists to search for similar patterns and relationships for other ecosystems and to find “Redfield-like” ratios to understand the balance of chemical elements in an ecological context (Sterner and Elser 2002; Cleveland and Liptzin 2007). In its broadest term, Redfield-like stoichiometry is achieved when biota have clearly defined (constrained) nutrient ratios. Conceptually this constrained biotic stoichiometry may imprint itself on the environment, when the flow of nutrients across the system boundaries (e.g. ocean surface, water/sediment boundary) are small relative to biota internal cycling rates (ingestion and egestion) combined with pathways of preferential losses of excess nutrients (Lenton and Watson 2000). The development of a Redfield-like ratio in a system is further hastened when biota can also preferentially acquire elements from external sources (e.g. biological N fixation). The combined outcome of in versus out (i.e. biogeochemical balancing) with sufficient time to reach equilibration/steady state ultimately produces a consistent nutrient stoichiometry (Lenton and Watson 2000; Sterner 2008).

While proportional (constrained) stoichiometric relationships between nutrients is the underpinning concept of the classic Redfield hypothesis as discussed above, relationships between C, N and P departed from isometric (proportional) relationships within most compartments (water column, floc, soil, and vegetation) in our study. Exceptions occur at specific locations within a FW notably between C and N in floc and soil compartments of FW 3 and FW 1, respectively (Table 3, Fig 8 and Fig 9). Much more prevalent was allometric (independent) scaling with slope values different from one (*β* ≠ 1). This deviation from an isometric relationship indicates the relative enrichment or depletion of one element relative to another resulting in lower or higher stoichiometric ratios (Fig 1). In most cases, the slope between nutrient concentrations were less than one (Table 2 and 3) indicating that the relative change of a more abundant element compared to the relative change to a less abundant element is muted. For example, an increase in P leads to comparatively smaller increases in N and C (Fig 3, 4 and 5).

**Figure 8.**
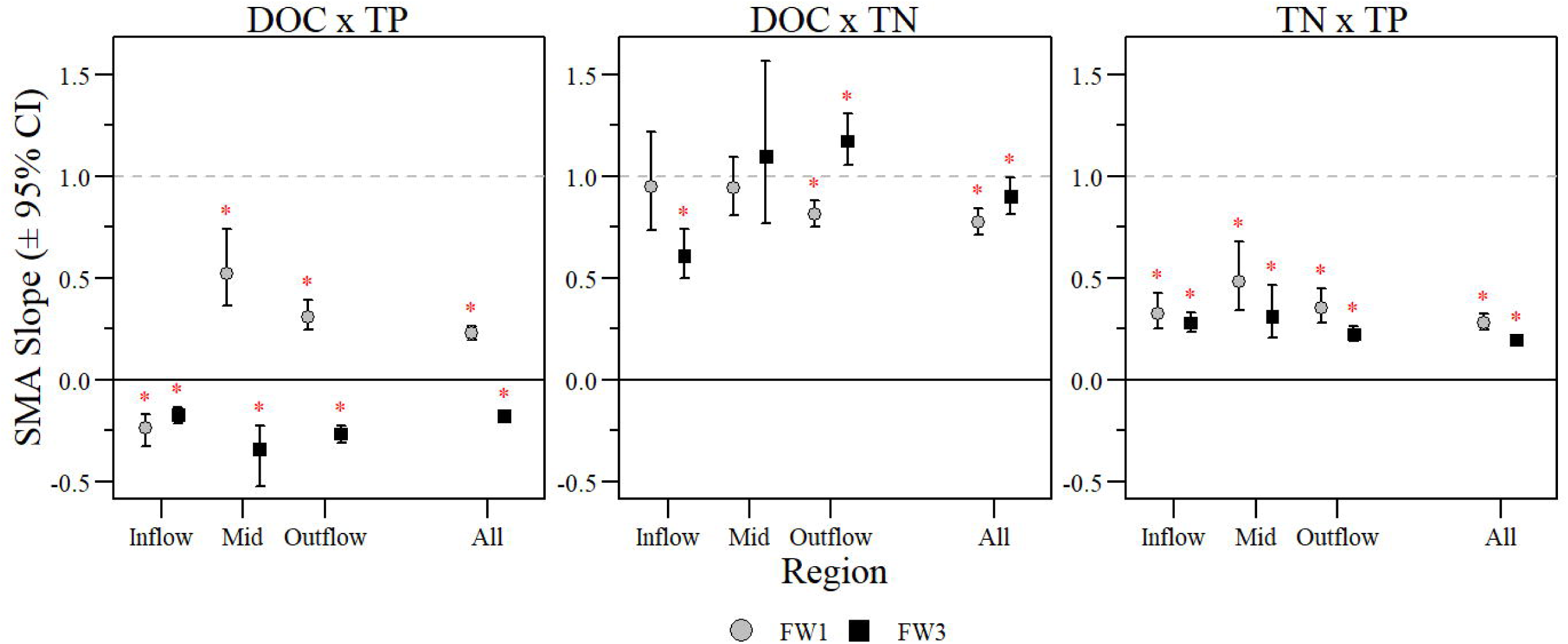
Surface water inflow, mid, outflow and overall standardized major axis slope values with ± 95% confidence interval for dissolved organic carbon (DOC), total phosphorus (TP) and total nitrogen (TN) comparisons. Slopes significantly different from one are identified with red asterisks above the upper 95% confidence interval bar.

**Figure 9.**
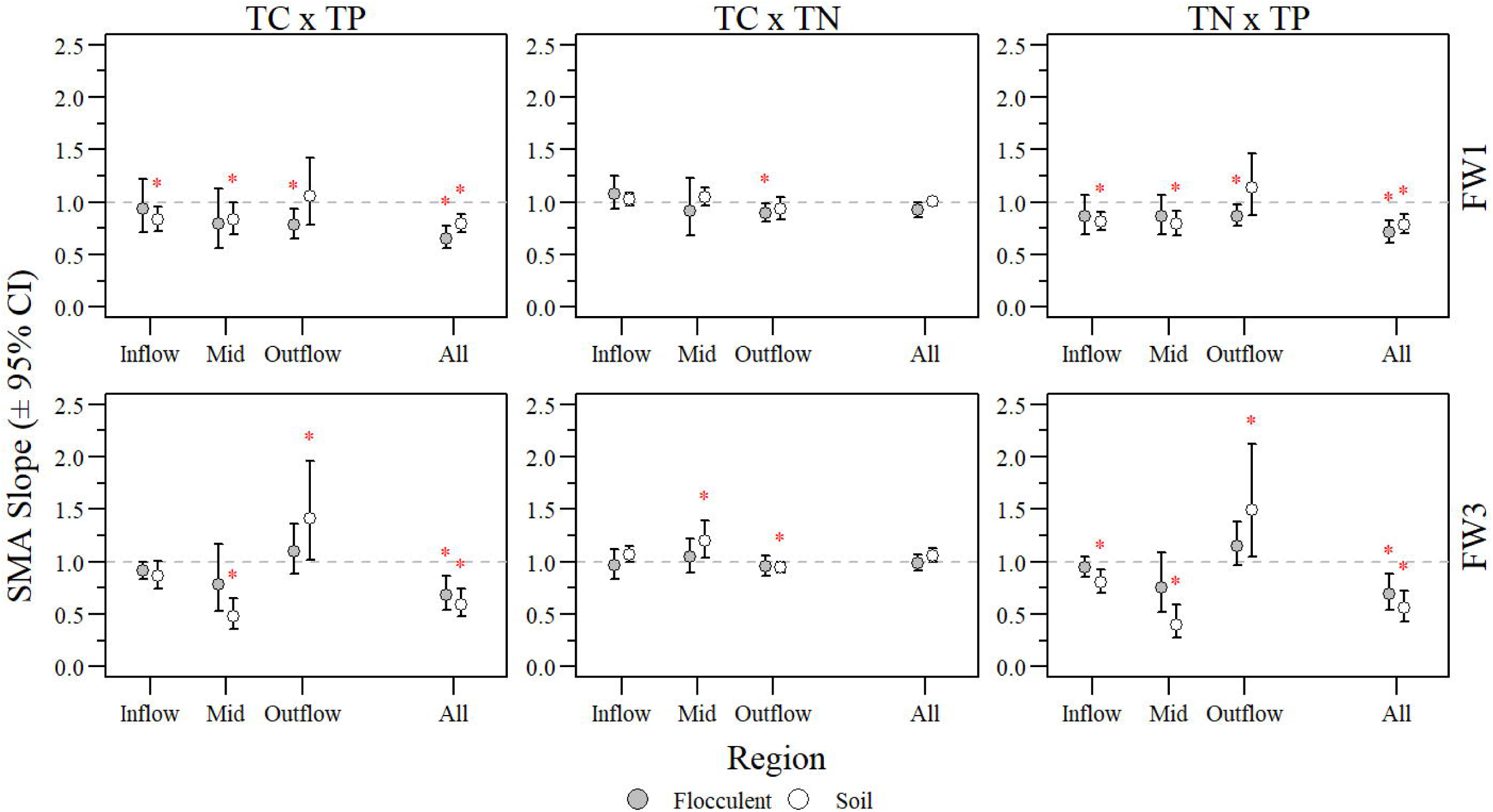
Flocculent and soil inflow, mid, outflow and overall standardized major axis slope values with ± 95% confidence interval for total carbon (TC), total phosphorus (TP) and total nitrogen (TN) comparisons. Slopes significantly different from one are identified with red asterisks above the upper 95% confidence interval bar.

In the water column, even under optimal nutrient use efficiency that is necessary for generating isometric scaling (Sterner et al. 2008), short residence time combined with large and variable external nutrient loading (Table S2 and Fig S2) may prevent adjustment to a potential locally consistent Redfield-like nutrient stoichiometry similar to observations found in tropical streams and lakes (They et al. 2017). The biogeochemical mosaic hypothesis addresses this cross-scale contrasts in nutrient dynamics recognizing that biogeochemical processes occur under contrasting conditions and spatially separated therefore occurring at different temporal and spatial scale thereby influencing (or hampering) the development of Redfield-like relationships across ecosystem compartments (Sterner et al. 2008). As observed in this study, inflow regions experience the highest nutrient load to the FWs and as the water moves along the FW total water column nutrients including inorganic nutrients are reduced to background concentrations, as per the design and intent of the system (Goforth 2007; Chen et al. 2015; UF-WBL 2017). Additionally, considerable allochthonous inputs of nutrients act as resource subsidy where they exert a strong influence on metabolism and material cycling and may be a critical component to deviations from the characteristic Redfield ratio (Hessen et al. 2003). Moreover, deviations from any Redfield-like ratio can have large consequences at larger spatial scales possibly to the point of system stress (Odum et al. 1979; Valett et al. 2008). Given these conditions, allometric (independent) scaling in the water column is expected and confirmed by our analysis (Table 2, Fig 3 and Fig 8). Differences in nutrient relationships may further be affected by nutrient movement (i.e. dispersion, advection, burial, bioturbation, etc.) from one compartment to another. In this study, nutrient ratios in vegetation, floc, and soil were different which allowed for stoichiometric mixing between compartments with differences between and along FWs (Table 3). For example, high net aquatic productivity occurs within the water column in the SAV dominated FW 3 (UF-WBL 2017), creating conditions for abiotic immobilization and deposition of nutrients (Juston et al. 2013; Zamorano et al. 2018). In contrast, primary productivity (low net aquatic productivity; UF-WBL 2017) occurs outside the water column in the EAV system, likely rendering the water column a dominantly heterotrophic system, as opposed to a predominantly autotrophic system in FW 3.

The relative position of the SMA regression lines in the log-log space can infer changes in potential limitations and how one nutrient relates to another. Dodds and Smith (2016) using a large regional dataset demonstrated that based on water column TN and TP concentrations, the range of values predicts N limitation of algal growth in some ecosystems and P limitation in others based on the relative position of values compared to the N:P Redfield Ratio of 16:1. The classic N:P Redfield ratio of 16:1 indicates a roughly balanced supply of N and P, while algae assemblages generally mirror this ratio under balanced growth conditions. In freshwater systems, a TN:TP molar ratio of 20:1 may be a better indicator of algal nutrient limitation than dissolved inorganic fractions of N and P (Guildford and Hecky 2000). Regardless of the metric, the Everglades STAs are strongly P-limited (Walker 1995; Juston and DeBusk 2006; Walker and Kadlec 2011; Chen et al. 2015) and for this study (Fig 3) where all surface water TN to TP values were above the P limitation threshold. Moreover, this study suggests that FW 3 is more P-limiting than FW 1 based on the relative position of the data (SMA regression) to the 20:1 or 16:1 balanced N to P thresholds (Fig 3). The resulting SMA slope is slightly (but significantly) higher in FW 1, implying lower use efficiencies for C and N in the SAV dominated FW 3. However, it is not clear whether these efficiencies are the main driver of shallow SMA slopes, given the large differences in biotic and abiotic nutrient pathways discussed above.

The absence of a Redfield-like relationship (i.e. constrained SMA slopes ≈ 1) in the floc and soil in our dataset differs from previous work that sought to explore relationships in grassland and forest soils (Cleveland and Liptzin 2007). Prior studies evaluating the nutrient stoichiometric relationships within the soil compartment compared nutrients on a molar mass per mass of soil ratio (i.e. mmol kg^−1^)(Cleveland and Liptzin 2007; Xu et al. 2013). Cleveland and Liptzin (2007) evaluated forested and grassland ecosystems which typically have higher relative mineral C that highly co-varied with other nutrients. This was also largely the case for Xu et al. (2013) but their study did also include natural wetland ecosystems in the synthesis of stoichiometric relationships. Evaluation of this wetland specific data compiled by Xu et al. (2013) indicated very little variation of C relative to other nutrients consistent with Fig S4 and Fig S5 in this study which evaluated stoichiometry based on concentrations (mmol kg^−1^). However, in wetlands, much of the soil mass consists of C. Therefore, an increase in C changes the mass, which itself is used to calculate concentrations. In other words, at high C concentrations, the SMA slope approaches zero, if a macronutrient is a substantial part of the overall soil mass. For this study, soil nutrient concentrations were expressed on a per area basis, instead of the usual normalization by mass or volume. We demonstrate the effect of concentration per area (concentration per area, mol m^−2^) vs. concentration per mass (mol kg^−1^) in Fig S1.

Floc can be viewed as the beginning of soil OM diagenesis in natural wetland ecosystems, being a mixture of leaf litter, other organic matter (i.e. bacterial cells, phytoplankton, algae, consumers, fecal material, etc.) in various states of decay and inorganic particles (clays and silts) (Droppo 2001; Noe et al. 2003; Neto et al. 2006). This matrix of biologic, chemical and even geologic material is thus expected to be stoichiometrically sandwiched between primary producers (i.e. vegetation and algae) on the one hand and soil on the other hand. Conceptually, primary producers exhibit relatively high C to nutrient ratio, which may even widen stoichiometric values in fresh litter because of plant’s re-translocation before tissue abscission (McGroddy et al. 2004). Active microbial pools immobilize (or retain) nutrients during early stages of decomposition, while C is respired for energy resulting in relative nutrient enrichment (Reddy and DeLaune 2008). Generally, the reactivity of floc and soils differ significantly with floc being much more reactive than soils as indicated by higher nutrient mineralization rates, nutrient content and microbial activity (Wright and Reddy 2001b; Neto et al. 2006). In this decomposition continuum it is expected that N relative to C increases during the transition from vegetation to floc to soil, while C:P ratios are expected to decline from vegetation to the floc layer, but then increase again as P mining becomes the dominant P redistribution mechanism in the soil. This decomposition, redistribution and nutrient utilization dynamics is apparent in stoichiometric relationships observed in this study (Fig 4, 5, 6 and 7), with the notable difference that C:N ratios did not further narrow in their transition from floc to soil.

Inspection of N to P stoichiometric relationships between floc and soils also revealed critical differences between FW1 (EAV) and FW3 (SAV) (Fig 6). In FW3, Floc TN:TP ratios in mid and outflow regions of the FW with much lower nutrient loading hover mostly around 60 molar ratio, while soil ratios are widely dispersed and vary between ~60 to ~180. In contrast, in FW 1, floc TN:TP ratios varied by more than a factor of 2, while soil ratios had a much smaller range. Both FWs had critically narrower TN:TP ratios at the inflow comparted to other locations in the FW. The stoichiometry of the midflow section veered more away from inflow characteristics in FW 3 than FW 1. The lack of a N:P ceiling in soil (but not in floc) in FW 3 could indicate that the soil compartment is continued to be depleted in P with decreasing nutrient load to the system (i.e. increasing distance downstream). The N:P pattern between floc and soil suggests that the depletion of P relative to N most likely due vegetation uptake or abiotic immobilization occurring primarily in floc in FW 1, whereas P depletion occurred mostly in soil in FW 3. This notion is also confirmed by the differences in N:P slopes of the SMA, where the slopes for floc is shallower than for soil in FW 1, but steeper in FW 3.

Aboveground vegetation SMA slopes also exhibited differences between FWs (i.e. between EAVs and SAVs). Nutrient stoichiometric relationships in plants reflect to some degrees the balance between demands of plant growth and nutrient supply rates from sediment and surrounding water in aquatic ecosystems (Frost et al. 2002). Nutrient and light availability are key controls on the chemical quality of plant materials and its interaction with the detrital pool (Evans-White and Halvorson 2017). The relative relationships of C to N or P are also relevant to the structural composition of the plant where higher C:nutrient values indicate reduced allocation to low-nutrient structural material (Chimney and Pietro 2006; De Deyn et al. 2008). Nutrient composition in plant tissues can be an important feature to identify ecological strategies of species relative to biogeochemical conditions (Tilman 1982). Generally, EAV species invest a significant quantity of C in their biomass, associated with structural components of different plant parts and generally have higher net primary production than SAV species (Reddy and DeLaune 2008; De Deyn et al. 2008). To some degree, SAV could also be light-limited especially when optical water quality degrades due to the suspension of particulate matter (Evans-White and Halvorson 2017; Zamorano et al. 2018). However, this seems not to be the case in FW3 (SAV) with SMA slope suggesting an above-linear increase in C relative to nutrients and not indicating a limitation for C acquisition. Perhaps with increased nutrient supply, investments shift to more structural tissues. In contrast, in FW1, relationships remained isometric except for N:P.

Overall, the core concepts of ecological stoichiometry include stoichiometric homeostasis, threshold elemental ratio and the growth rate hypothesis which lay out the rules for differential nutrient demand, nutrient recycling, and nutrient transfer from one compartment to the next (Frost et al. 2002; Sterner and Elser 2002; Van de Waal et al. 2018). Within this framework, comparison of nutrient stoichiometry (C:N:P ratios) under differential nutrient supplies and across different systems provides insight into the nature of differential nutrient cycling rates and organization of material, and in particular also nutrient status within the ecosystem in question. The analysis here confirms earlier work that suggests primarily P limitation (UF-WBL 2017) where increasing P supply alleviates a deficit of this nutrient relative to others. This allometric relationship is pervasive in most compartments, especially in FW3 (SAV). The analysis from inflow to outflow shows a slightly more nuanced picture as the expectation is for increasing P limitation with distance downstream and thus increasing SMA slopes with P from inflow to outflow, yet this is more often refuted than confirmed for most compartments. However, drawing firm conclusions from SMA’s along the FW may be limited because of a possible limiting sample sizes and limited variation of nutrients at specific locations

Regarding constrained stoichiometry (allometry vs. isometry), an objective of this work was to determine whether the relationships are predictable (i.e. *coupled*), with sufficient variance explained in the SMA relationship. Thus, our definition of coupled accepts both isometric and allometric scaling (Fig 1). We find that the log-log regressions explain much less variance in the water column compared to other compartments, to a point that is borderline decoupled by our definition. A possible mechanism to cause decoupled or unconstrained stoichiometric relationships in a system would be an external nutrient load that is variable and overwhelms the local system. High external loads with an “unbalanced” stoichiometry, especially in the case of stormwater run-off where waters can have a disproportionate amount of N or P ultimately disrupting local nutrient cycling, and subsequently local stoichiometric ratios. If such load is large and variable, the system is then in a pervasive state of disequilibrium, leaving the water column in a borderline decoupled state (low R^2^) with variable and largely unpredictable stoichiometry. Ultimately, allochthonous inputs have the potential to alter endogenous processes such as nutrient cycling and decomposition of OM. In this study, nutrient load was variable across sampling events occurring during specific flow events with large fluctuation in input of new nutrients (Fig S2). This may be a major reason that stoichiometric relationships in the water column appear to be close to a decoupled state (i.e. R^2^ < 0.25). However, on the longer timescales for which vegetation, floc and soil operate, stoichiometry appears to become more coupled and thus predictable (high R^2^). It is interesting, though, that there is no evidence of stoichiometry becoming more coupled along the FW, suggesting that the mechanisms of preferential removal or selective fixation cannot restore the stoichiometric imbalance in the water column over the time it takes to move from inflow to outflow.

While not explicitly studied here, differences in uptake rate and nutrient form (i.e. dissolved versus particulate) can influence stoichiometric relationships, especially when stoichiometry is analyzed simply based on the water column nutrient content without considering the biotic component and a nutrient source. The prevailing paradigm of the Redfield ratio is consistency in both, the mineral nutrients and the organic matter which applies well to oceanic systems. Meanwhile in lake ecosystems, spatial and temporal variation in nutrient supply, overall higher seston nutrient content as well as contrasting system configurations and hydrodynamics when compared to oceanic systems contribute to more overall variability in composition and stoichiometry (Sterner et al. 2008). Generally, the water column of FW1 and FW3 showed some characteristics of a lake ecosystem more so than a marine ecosystem exhibiting a high degree of variability in nutrient stoichiometry. The biogeochemistry of treatment wetlands characterized here create a systematically decreasing load along the FW in which sedimentation of organic particulates and transport of dissolved nutrients may lead to preferential stoichiometry of retention in different compartments developing a strong spatial chemical gradient (Engle and Melack 1993; Sánchez-Carrillo et al. 2001; Angeler et al. 2007). This is evident in our study with slope values along FWs being highly variable with generally higher variability at the inflow regions especially with respect to DOC to TN comparisons (Fig 8).

### Conclusion and Further Research

Prior studies of stoichiometry suggest that the relationship between C and nutrients is tightly constrained and C:N:P stoichiometric relationships are relatively constrained and consistent with elemental composition of dominant phototrophs (i.e. algae and phytoplankton) in the water column and microbial biomass in soil (Redfield 1958; Elser et al. 2000, 2007; Cleveland and Liptzin 2007; Sterner et al. 2008; Xu et al. 2013). At a finer scale, as exemplified in this study nutrient stoichiometric relationships within treatment wetlands evaluated in this study were unconstrained through processes influenced by high loading resulting in nutrient enrichment causing disruption of biotic-feedback loops, variable mineralization/immobilization rates, selective removal of water column constituents via biotic uptake or, physical settling, and/or active mining for limiting nutrients in soils. As a consequence, stoichiometric relationships across different ecosystem compartments varied along nutrient gradients and ecosystem types. At the onset of this study two hypotheses were suggested. The first hypothesis which stated that stoichiometric relationships will differ between FWs was supported as the comparison of SMA slopes was significantly different for most stoichiometric relationships except for TC by TP and TN by TP in the floc compartment and for TN by TP in live AGB. The second hypothesis that stoichiometric relationships will change along FWs was supported with most FW regional (i.e. Inflow, Mid and Outflow) SMA slopes being significantly different within FWs but not consistently between FWs. Moreover, based on FW regional SMA model results stoichiometric scaling within these regions and the degree of coupling was highly variable along each FW with respect to C, N and P stoichiometry (Table 2 and 3).

Evaluation of stoichiometric relationships within ecosystem compartments provides a greater understanding of how nutrients and OM cycle through a system relative to one another (i.e. relative enrichment or depletion) and the degree of process coupling such as the diametrically opposed relationships between DOC and TP in the water column along each FW. Scaling properties of nutrients, as indicated by power law slopes in a stoichiometric framework, provide an understanding of fundamental relationships and processes in natural systems, and variability in these relationships can offer insight into underlying biogeochemical mechanisms (Brown et al. 2002; Marquet et al. 2005; Wymore *Unpublished Data*). Divergent stoichiometric relationships of C, N and P from a proportional scaling model (i.e. isometric; Fig 1) suggest how nutrients are retained within the ecosystem could be used to understand treatment wetland performance and expectations. In the context of the Everglades STAs, optimization for P-retention must go beyond a focus on P and P-forms (i.e. organic versus inorganic) but also focus on other constituents including C and N. Building from this work, future studies should address the potential for preferential removal and utilization of nutrients from different substrates and organisms (uptake and mining in macrophytes, immobilization of nutrients) to further understand ecosystem nutrient uptake and retention.

## Acknowledgements

We would like to thank SFWMD and UF Wetland Biogeochemistry Laboratory staff members for providing the data used in this analysis. We would also like to thank Mark Brenner, Sue Newman, K. Ramesh Reddy, Odi Villapando, Delia Ivanoff and the anonymous peer reviewer(s) and editor(s) for their efforts and constructive review of this manuscript.

## Authors’ Contributions

PJ performed data analyses including necessary calculations and statistical analyses and wrote the manuscript. SG and ALW contributed to data interpretation and writing. RKB and TZO were involved with data collection and writing. JK, MP and JD were involved in data collection. All authors read and approved the final manuscript.

## References

Angeler DG, Sánchez-Carrillo S, Rodrigo MA, et al (2007) Does seston size structure reflect fish-mediated effects on water quality in a degraded semiarid wetland? Environ Monit Assess 125:9–17. doi: 10.1007/s10661-006-9234-5

Bhomia RK, Reddy KR (2018) Influence of Vegetation on Long-term Phosphorus Sequestration in Subtropical Treatment Wetlands. J Environ Qual 47:361. doi: 10.2134/jeq2017.07.0272

Brown JH, Gupta VK, Li B-L, et al (2002) The fractal nature of nature: power laws, ecological complexity and biodiversity. Philos Trans R Soc B Biol Sci 357:619–626. doi: 10.1098/rstb.2001.0993

Bruland GL, Osborne TZ, Reddy KR, et al (2007) Recent Changes in Soil Total Phosphorus in the Everglades: Water Conservation Area 3. Environ Monit Assess 129:379–395. doi: 10.1007/s10661-006-9371-x

Chen H, Ivanoff D, Pietro K (2015) Long-term phosphorus removal in the Everglades stormwater treatment areas of South Florida in the United States. Ecol Eng 79:158–168. doi: 10.1016/j.ecoleng.2014.12.012

Chimney M (2017) Performance of the Everglades Stormwater Treatment Areas. In: 2017 South Florida Environmental Report. South Florida Water Management District, West Palm Beach, FL

Chimney MJ, Pietro KC (2006) Decomposition of macrophyte litter in a subtropical constructed wetland in south Florida (USA). Ecol Eng 27:301–321. doi: 10.1016/j.ecoleng.2006.05.016

Cleveland CC, Liptzin D (2007) C:N:P stoichiometry in soil: is there a “Redfield ratio” for the microbial biomass? Biogeochemistry 85:235–252. doi: 10.1007/s10533-007-9132-0

Collins SM, Oliver SK, Lapierre J-F, et al (2017) Lake nutrient stoichiometry is less predictable than nutrient concentrations at regional and sub-continental scales. Ecol Appl 27:1529–1540. doi: 10.1002/eap.1545

Corstanje R, Grafius DR, Zawadzka J, et al (2016) A datamining approach to identifying spatial patterns of phosphorus forms in the Stormwater Treatment Areas in the Everglades, US. Ecol Eng 97:567–576. doi: 10.1016/j.ecoleng.2016.10.003

Davis SM (1991) Growth, decomposition, and nutrient retention of Cladium jamaicense Crantz and Typha domingensis Pers. in the Florida Everglades. Aquat Bot 40:203–224. doi: 10.1016/0304-3770(91)90059-E

De Deyn GB, Cornelissen JHC, Bardgett RD (2008) Plant functional traits and soil carbon sequestration in contrasting biomes. Ecol Lett 11:516–531. doi: 10.1111/j.1461-0248.2008.01164.x

Dodds W, Smith V (2016) Nitrogen, phosphorus, and eutrophication in streams. Inland Waters 6:155–164. doi: 10.5268/IW-6.2.909

Dodds WK (2003) Misuse of inorganic N and soluble reactive P concentrations to indicate nutrient status of surface waters. J North Am Benthol Soc 22:171–181

Dodds WK, Smith VH, Lohman K (2002) Nitrogen and phosphorus relationships to benthic algal biomass in temperate streams. Can J Fish Aquat Sci 59:865–874. doi: 10.1139/f02-063

Dombrowski J, Powers M, Zamorano M, et al (2018) Appendix 5B-5: Submerged Aquatic Veetation Coverage in the Stormwater Treatment Areas Based on Ground Surveys. In: South Florida Environmental Report. South Florida Water Management District, West Palm Beach, FL

Droppo IG (2001) Rethinking what constitutes suspended sediment. Hydrol Process 15:1551–1564. doi: 10.1002/hyp.228

Elser JJ, Andersen T, Baron JS, et al (2009) Shifts in Lake N:P Stoichiometry and Nutrient Limitation Driven by Atmospheric Nitrogen Deposition. Science 326:835–837. doi: 10.1126/science.1176199

Elser JJ, Bracken MES, Cleland EE, et al (2007) Global analysis of nitrogen and phosphorus limitation of primary producers in freshwater, marine and terrestrial ecosystems. Ecol Lett 10:1135–1142. doi: 10.1111/j.1461-0248.2007.01113.x

Elser JJ, Fagan WF, Denno RF, et al (2000) Nutritional constraints in terrestrial and freshwater food webs. Nature 408:578

Engle DL, Melack JM (1993) Consequences of riverine flooding for seston and theperiphyton of floating meadows in an Amazon floodplain lake. Limnol Oceanogr 38:1500–1520. doi: 10.4319/lo.1993.38.7.1500

Evans-White MA, Halvorson HM (2017) Comparing the Ecological Stoichiometry in Green and Brown Food Webs – A Review and Meta-analysis of Freshwater Food Webs. Front Microbiol 8:. doi: 10.3389/fmicb.2017.01184

Frost PC, Stelzer RS, Lamberti GA, Elser JJ (2002) Ecological stoichiometry of trophic interactions in the benthos: understanding the role of C:N:P ratios in lentic and lotic habitats. J North Am Benthol Soc 21:515–528. doi: 10.2307/1468427

Geider R, La Roche J (2002) Redfield revisited: variability of C:N:P in marine microalgae and its biochemical basis. Eur J Phycol 37:1–17. doi: 10.1017/S0967026201003456

Goforth G (2007) Updated STA Phosphorus Modeling for the 2010 Planning Period. South Florida Water Management District, West Palm Beach, FL

Guildford SJ, Hecky RE (2000) Total nitrogen, total phosphorus, and nutrient limitation in lakes and oceans: Is there a common relationship? Limnol Oceanogr 45:1213–1223. doi: 10.4319/lo.2000.45.6.1213

Helton AM, Ardón M, Bernhardt ES (2015) Thermodynamic constraints on the utility of ecological stoichiometry for explaining global biogeochemical patterns. Ecol Lett 18:1049–1056. doi: 10.1111/ele.12487

Hessen DO, Andersen T, Brettum P, Faafeng BA (2003) Phytoplankton contribution to sestonic mass and elemental ratios in lakes: Implications for zooplankton nutrition. Limnol Oceanogr 48:1289–1296. doi: 10.4319/lo.2003.48.3.1289

Howard-Williams C (1985) Cycling and retention of nitrogen and phosphorus in wetlands: a theoretical and applied perspective. Freshw Biol 15:391–431. doi: 10.1111/j.1365-2427.1985.tb00212.x

Julian P, Gerber S, Wright AL, et al (2017) Carbon pool trends and dynamics within a subtropical peatland during long-term restoration. Ecol Process 6:43–57. doi: 10.1186/s13717-017-0110-8

Juston J, DeBusk TA (2006) Phosphorus mass load and outflow concentration relationships in stormwater treatment areas for Everglades restoration. Ecol Eng 26:206–223. doi: 10.1016/j.ecoleng.2005.09.011

Juston JM, DeBusk TA (2011) Evidence and implications of the background phosphorus concentration of submerged aquatic vegetation wetlands in Stormwater Treatment Areas for Everglades restoration. Water Resour Res 47:. doi: 10.1029/2010WR009294

Juston JM, DeBusk TA, Grace KA, Jackson SD (2013) A model of phosphorus cycling to explore the role of biomass turnover in submerged aquatic vegetation wetlands for Everglades restoration. Ecol Model 251:135–149. doi: 10.1016/j.ecolmodel.2012.12.001

Kadlec RH, Wallace SD (2009) Treatment wetlands. CRC Press, Boca Raton, FL

Lenton TM, Watson AJ (2000) Redfield revisited: 1. Regulation of nitrate, phosphate, and oxygen in the ocean. Glob Biogeochem Cycles 14:225–248

Marquet PA, Quinones RA, Abadas S, et al (2005) Scaling and power-laws in ecological systems. J Exp Biol 208:1749–1769. doi: 10.1242/jeb.01588

McGroddy ME, Daufresne T, Hedin LO (2004) Scaling of C:N:P stoichiometry in forests worldwide: implications of terrestrial redfield-type ratios. Ecology 85:2390–2401. doi: 10.1890/03-0351

Neto RR, Mead RN, Louda JW, Jaffé R (2006) Organic Biogeochemistry of Detrital Flocculent Material (Floc) in a Subtropical, Coastal Wetland. Biogeochemistry 77:283–304. doi: 10.1007/s10533-005-5042-1

Newman S, McCormick PV, Miao SL, et al (2004) The effect of phosphorus enrichment on the nutrient status of a northern Everglades slough. Wetl Ecol Manag 12:63–79. doi: 10.1023/B:WETL.0000021664.32137.dd

Newman S, Osborne TZ, Hagerthey SE, et al (2017) Drivers of landscape evolution: multiple regimes and their influence on carbon sequestration in a sub-tropical peatland. Ecol Monogr 87:578–599. doi: 10.1002/ecm.1269

Noe GB, Scinto LJ, Taylor J, et al (2003) Phosphorus cycling and partitioning in an oligotrophic Everglades wetland ecosystem: a radioisotope tracing study. Freshw Biol 48:1993–2008. doi: 10.1046/j.1365-2427.2003.01143.x

Odum EP, Finn JT, Franz EH (1979) Perturbation Theory and the Subsidy-Stress Gradient. BioScience 29:349–352. doi: 10.2307/1307690

Osborne TZ, Bruland GL, Newman S, et al (2011a) Spatial distributions and eco-partitioning of soil biogeochemical properties in the Everglades National Park. Environ Monit Assess 183:395–408. doi: 10.1007/s10661-011-1928-7

Osborne TZ, Newman S, Scheidt DJ, et al (2011b) Landscape Patterns of Significant Soil Nutrients and Contaminants in the Greater Everglades Ecosystem: Past, Present, and Future. Crit Rev Environ Sci Technol 41:121–148. doi: 10.1080/10643389.2010.530930

Reddy KR, DeLaune RD (2008) Biogeochemistry of wetlands: science and applications. CRC Press, Boca Raton, FL

Reddy KR, Kadlec RH, Flaig E, Gale PM (1999) Phosphorus Retention in Streams and Wetlands: A Review. Crit Rev Environ Sci Technol 29:83–146. doi: 10.1080/10643389991259182

Redfield AC (1934) On the proportions of organic derivations in seawater and their relation to the composition of plankton (reprint). Benchmark Pap Ecol 1:

Redfield AC (1958) The biological control of chemical factors in the environment. Am Sci 46:230A–221

Sánchez-Carrillo S, Álvarez-Cobelas M, Angeler DG (2001) Sedimentation in the semi-arid freshwater wetland las tablas de Daimiel (Spain). Wetlands 21:112–124. doi: 10.1672/0277-5212(2001)021[0112:SITSAF]2.0.CO;2

SFWMD (2012) Restoration strategies regional water quality plan. South Florida Water Management District, West Palm Beach, FL

Sterner RW (2008) On the Phosphorus Limitation Paradigm for Lakes. Int Rev Hydrobiol 93:433–445. doi: 10.1002/iroh.200811068

Sterner RW, Andersen T, Elser JJ, et al (2008) Scale-dependent carbon: nitrogen: phosphorus seston stoichiometry in marine and freshwaters. Limnol Oceanogr 53:1169

Sterner RW, Elser JJ (2002) Ecological Stoichiometry: The Biology of Elements from Molecules to the Biosphere. Princeton University Press

They NH, Amado AM, Cotner JB (2017) Redfield Ratios in Inland Waters: Higher Biological Control of C:N:P Ratios in Tropical Semi-arid High Water Residence Time Lakes. Front Microbiol 8:. doi: 10.3389/fmicb.2017.01505

Tilman D (1982) Resource Competition and Community Structure. Princeton University Press

UF-WBL (2017) Evaluation of Soil Biogeochemical Properties Influencing Phosphorus Flux in the Everglades Stormwater Treatment areas: 2016-2017 Annual Report. University of Florida, Gainesville, FL

Valett HM, Thomas SA, Mulholland PJ, et al (2008) Endogenous and Exogenous Control of Ecosystem Function: N Cycling in Headwater Streams. Ecology 89:3515–3527. doi: 10.1890/07-1003.1

Van de Waal DB, Elser JJ, Martiny AC, et al (2018) Editorial: Progress in Ecological Stoichiometry. Front Microbiol 9:. doi: 10.3389/fmicb.2018.01957

Walker WW (1995) Design Basis for Everglades Stormwater Treatment Areas. JAWRA J Am Water Resour Assoc 31:671–685. doi: 10.1111/j.1752-1688.1995.tb03393.x

Walker WW, Kadlec RH (2011) Modeling Phosphorus Dynamics in Everglades Wetlands and Stormwater Treatment Areas. Crit Rev Environ Sci Technol 41:430–446. doi: 10.1080/10643389.2010.531225

Warton DI, Duursma RA, Falster DS, Taskinen S (2012) smatr 3-an R package for estimation and inference about allometric lines: The smatr 3-an R package. Methods Ecol Evol 3:257–259. doi: 10.1111/j.2041-210X.2011.00153.x

Warton DI, Weber NC (2002) Common Slope Tests for Bivariate Errors-in-Variables Models. Biom J 44:161–174. doi: 10.1002/1521-4036(200203)44:2<161::AID-BIMJ161>3.0.CO;2-N

Warton DI, Wright IJ, Falster DS, Westoby M (2006) Bivariate line-fitting methods for allometry. Biol Rev 81:259–291. doi: 10.1017/S1464793106007007

Wright AL, Reddy KR (2001a) Phosphorus loading effects on extracellular enzyme activity in Everglades wetland soils. Soil Sci Soc Am J 65:588–595

Wright AL, Reddy KR (2001b) Heterotrophic Microbial Activity in Northern Everglades Wetland Soils. Soil Sci Soc Am J 65:1856. doi: 10.2136/sssaj2001.1856

Wymore AS, Brereton RL, Ibarra DE, et al (2017) Critical zone structure controls concentration-discharge relationships and solute generation in forested tropical montane watersheds. Water Resour Res 53:6279–6295. doi: 10.1002/2016WR020016

Xu X, Thornton PE, Post WM (2013) A global analysis of soil microbial biomass carbon, nitrogen and phosphorus in terrestrial ecosystems. Glob Ecol Biogeogr 22:737–749. doi: 10.1111/geb.12029

Zamorano MF, Bhomia RK, Chimney MJ, Ivanoff D (2018) Spatiotemporal changes in soil phosphorus characteristics in a submerged aquatic vegetation-dominated treatment wetland. J Environ Manage 228:363–372. doi: 10.1016/j.jenvman.2018.09.032

