## Supplemental Material for "Evaluation of nutrient stoichiometric relationships amongst ecosystem compartments of a subtropical treatment wetland. Do we have “Redfield Wetlands”?"

Table S1. Summary of parameters, matrices, analytical methods and minimum detection limit (MDL) used for this study. Additional parameters were collected as part of the larger study (Reddy 2017) but not used in this study. All analytical methods are consistent with Florida Department of Environmental Protection or U.S. Environmental Protection Agency Standard Operating Procedures and methods.

| **Matrix** | **Parameter** | **Abbreviation** | **Analytical Method** | **Minimum Detection Limit** | **Method** |
| --- | --- | --- | --- | --- | --- |
| Surface Water | Total Phosphorus | TP | SM4500PF | 2 µg P L^-1^ | Clesceri et al. (1998) |
|  | Total Nitrogen | TN | SM4500NC | 0.02 mg N L^-1^ | Clesceri et al. (1998) |
|  | Dissolved Organic Carbon | DOC | SM5310B | 0.8 mg C L^-1^ | Clesceri et al. (1998) |
| Soil and Vegetation | Loss-on-ignition^1,2^ | LOI | Calculation^2^ | 1.0 % | **---** |
|  | Total Phosphorus | TP | SM4500PF | 16 mg P kg^-1^ | Clesceri et al. (1998) |
|  | Total Nitrogen | TN | SFWMD 3200 | 2 g N kg^-1^ | SFWMD (2015) |
|  | Total Carbon | TC | SFWMD 3200 | 2 g C kg^-1^ | SFWMD (2015) |

^1^ Loss-on-ignition was assessed for soil components only.

^2^ Loss-on-ignition was calculated from the difference between 100% and percent ash determined by the analytical method identified as SFWMD 1610 (SFWMD 2015)**.**

Table S2**.** Event characteristics including duration, hydraulic and phosphorus loading rates (HLR and PLR, respectively) and median detection time of the five flow events for STA-2 flow-ways (FWs) 1 and 3.

| **STA Flow-way** | **Event** | **Start - End Date** | **Duration (Days)** | **HLR**  **(cm d⁻¹)** | **PLR**  **(mg m⁻² d⁻¹)** | **Median**  **Detention**  **Time (d)** |
| --- | --- | --- | --- | --- | --- | --- |
| FW 1 | 1 | Aug 10 - Sep 14, 2015 | 35 | 0.56 ± 0.12 | 0.46 ± 0.10 | 35.7 |
| FW 1 | 2 | Oct 20 - Nov 29, 2015 | 40 | 1.20 ± 0.25 | 0.37 ± 0.08 | 27.1 |
| FW 3 | 3 | Feb 22 - Apr 11, 2016 | 49 | 3.17 ± 0.63 | 1.45 ± 0.30 | 2.3 |
| FW 3 | 4 | Jun 27 - Aug 29, 2016 | 63 | 2.00 ± 0.24 | 1.16 ± 0.16 | 16.0 |
| FW 3 | 5 | Oct 12 - Nov 22, 2016 | 41 | 5.88 ± 0.84 | 4.33 ± 0.68 | 5.2 |
| FW 1 | 6 | May 29 - Jul 31, 2017 | 63 | 3.84 ± 0.71 | 7.10 ± 1.39 | 25.0 |

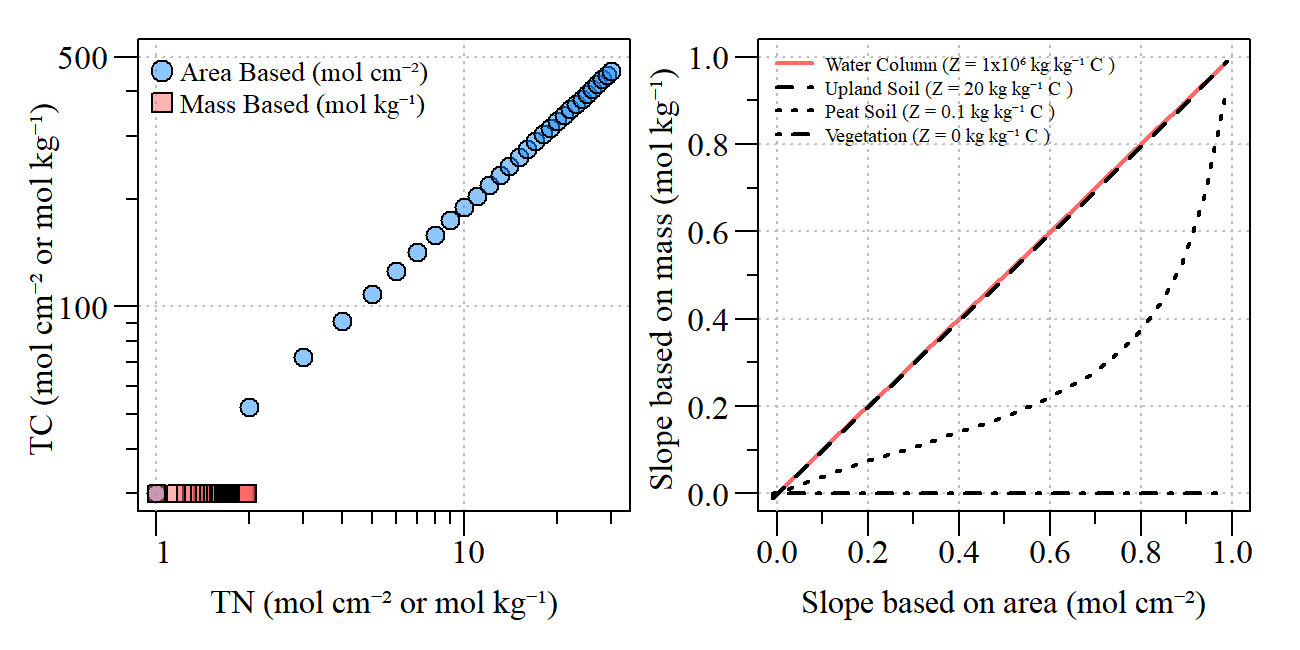

Figure S1. Effect of large concentration of carbon in area or mass on SMA regression. We assumed that the mass in a sample consists of C/0.45 + Z, where 0.45 is the concentration of C in biomass, and Z is material other than organics (minerals, water) on a area basis. The left figure assumes Z=0, as it may occur in vegetation, where carbon and a nutrient (N) scale allometrically with a SMA slope of 0.8. The slope becomes zero if normalized on a per mass basis (note, we normalized the values so that they are identical at log(N)=0). We then accounted for different amounts of non-organics (varying Z, bottom). Sufficient high Z lead to convergence of SMA slopes, but differences occur with decreasing Z, and become substantial for peat soils as analyzed in this study.

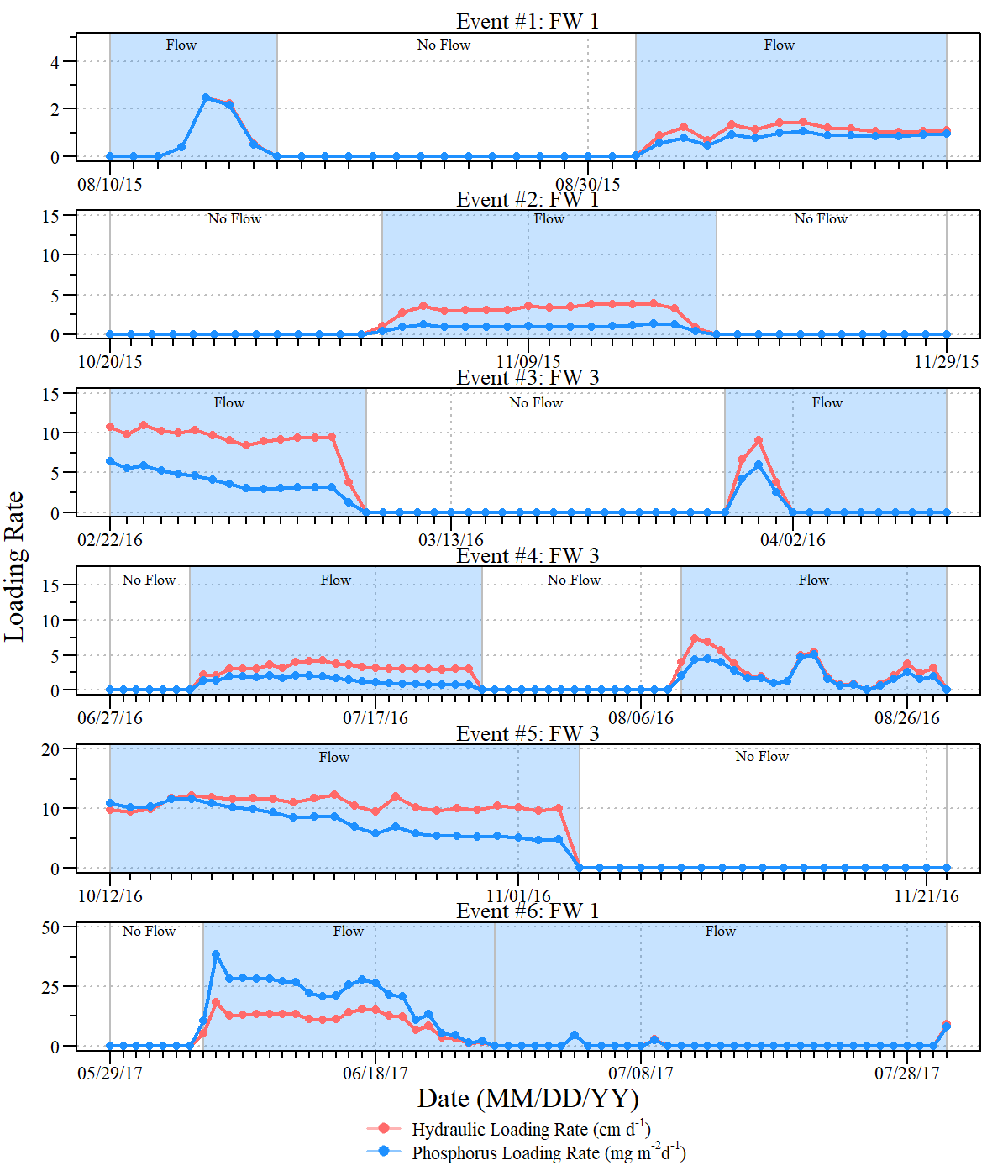

Fig S2. Hydrologic and Phosphorus loading rates (HLR and PLR, respectively) for the six flow events within flow-ways (FWs) 1 and 3 of Stormwater Treatment Area-2 between August 10^th^, 2015 and July 31^st^, 2017.

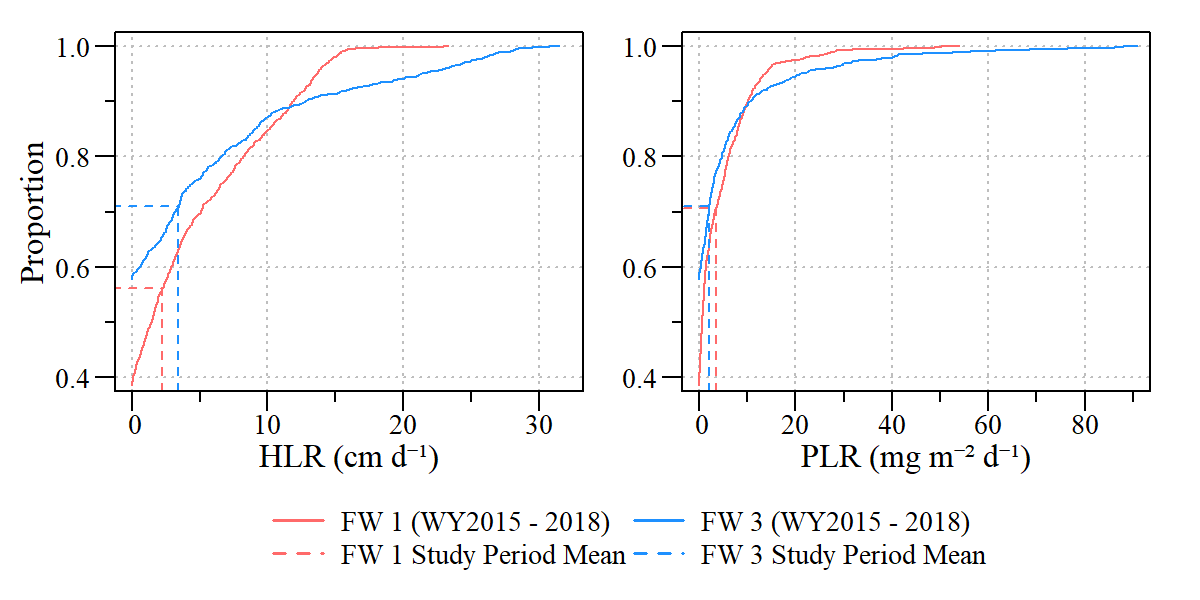

Fig S3. Cumulative distribution plots of hydraulic (left) and phosphorus (right) loading rates for flow-ways 1 and 2 (FW1 and FW2, respectively) for data collected between May 1^st^ 2014 and April 30^th^ 2018 (solid lines) relative to experiment flow period mean values (dashed values) for each flow-way.

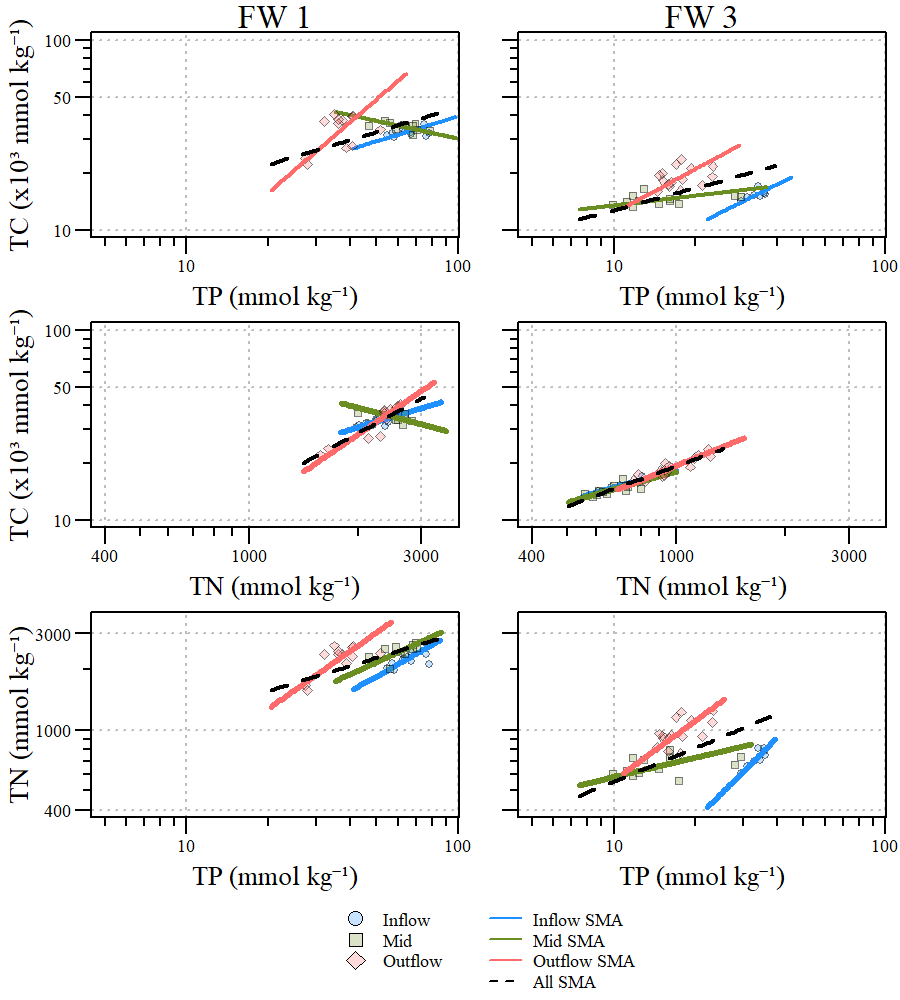

Figure S4. Stoichiometric relationships between total carbon (TC), total phosphorus (TP) and total nitrogen (TN) in floc ecosystem compartment for Stormwater Treatment Area 2, flow-ways (FWs) 1 and 3. Inflow, mid, outflow and overall standardized major axis (SMA) regressions indicated by lines through the data. Values can be converted to mass per volume (i.e. milligram per kilogram) concentration by multiplying each value by its respective conversion factor (C = 12.01; N = 14.00; P = 30.97).

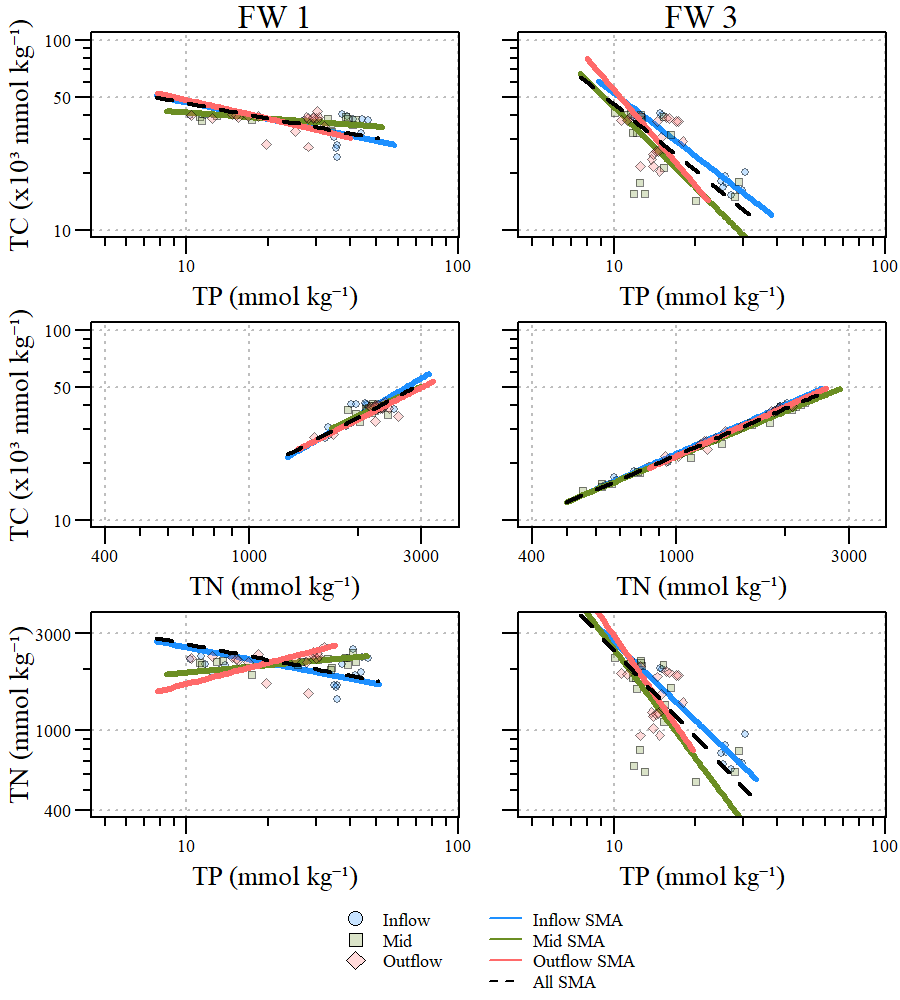

Figure S5. Stoichiometric relationships between total carbon (TC), total phosphorus (TP) and total nitrogen (TN) in soil ecosystem compartment for Stormwater Treatment Area 2, flow-ways (FWs) 1 and 3. Inflow, mid, outflow and overall standardized major axis (SMA) regressions indicated by lines through the data. Values can be converted to mass per volume (i.e. milligram per kilogram) concentration by multiplying each value by its respective conversion factor (C = 12.01; N = 14.00; P = 30.97).
